# Molecular basis of VEGFR1 autoinhibition at the plasma membrane

**DOI:** 10.1101/2023.06.08.544162

**Authors:** Manas Pratim Chakraborty, Diptatanu Das, Purav Mondal, Pragya Kaul, Soumi Bhattacharyya, Prosad Kumar Das, Rahul Das

## Abstract

Ligand-independent activation of VEGFR is a hallmark in diabetes and several cancers. Like most RTKs, the VEGFR2, the primary VEGF receptor, is activated spontaneously at higher receptor concentrations. An exception is VEGFR1, which remains constitutively inactive in the basal state. Ligand stimulation transiently phosphorylates VEGFR1 and induces weak kinase activation in endothelial cells. Recent studies, however, suggest that VEGFR1 signaling is indispensable in regulating various physiological or pathological events, which is puzzling. Why VEGFR1 is differentially regulated is an open question. Here we elucidate a mechanism of juxtamembrane inhibition that shifts the equilibrium more to the inactive state, rendering VEGFR1 an inefficient kinase. Our data suggest that a combination of tyrosine phosphatase activity and JM inhibition suppress the basal phosphorylation of VEGFR1. We conclude that a subtle imbalance in phosphatase activation or removing juxtamembrane inhibition is sufficient to induce basal activation of VEGFR1 and remodel tyrosine phosphorylation to be sustained.

## Introduction

The vascular endothelial growth factor receptors (VEGFR) are the key regulator of normal physiological and pathological angiogenesis and vasculogenesis (1,2). The VEGFR family comprises three receptor tyrosine kinases (RTK): VEGFR1, VEGFR2, and VEGFR3. Among them, VEGFR1 is an elusive family member. Even after three decades of its discovery, the function and regulation of VEGFR1 remain poorly understood (3–5). VEGFR2 is the primary receptor for VEGFs. It regulates diverse cellular functions, including blood vessel development during embryogenesis, hematopoiesis, and tumor angiogenesis (1,6). During embryonic development, VEGFR1 acts as a decoy receptor. VEGFR1 negatively regulates the VEGFR2 signaling by sequestering excess VEGF-A, preventing over activation of VEGFR2 (5,7,8) Compared to VEGFR2, VEGFR1 binds to its ligand VEGF-A with a ten-fold stronger affinity (9,10). Yet, the ligand binding induces only a weak kinase activation in VEGFR1 and does not generate subsequent downstream signaling in endothelial cells, vascular smooth muscle, or fibroblast cells (4,8,11,12). Although VEGFR1 and VEGFR2 share a high degree of sequence and structural homology, it is unclear why the two RTK are differently regulated.

VEGFR1 and VEGFR2 share similar structural architecture, comprising an extracellular ligand-binding domain (ECD) made up of seven immunoglobulin-like subdomains (D1 to D7), a single-passed transmembrane (TM) segment, a cytosolic juxtamembrane (JM) segment tethered to a kinase domain (KD) followed by a C-terminal tail (Figure 1A) (1,2,13). In the unligated state, the receptor exists predominantly as a monomer (Figure 1B) (14,15), and the KD adopts a PDGFR-like JM-in inactive conformation (16–19). The VEGFR1 and VEGFR2 are activated by a common bivalent ligand (VEGF-A), leading to a ligand-dependent dimerization of the ECD (Figure 1B)(20,21). The ECD dimerization rearranges the TM segment (22), removes the JM-inhibition (to JM-out conformation), and brings two adjacent KD in close proximity allowing autophosphorylation of multiple tyrosine residues in the C-terminal tail (Figure 1B)(16,18,20). The phosphotyrosine residues then function as a docking site for assembling downstream signaling modules. Structural analysis of the KD suggests that VEGFR1 is not a pseudokinase. All the regulatory motifs (R-spine and C-spine) and the catalytic residues are conserved in the VEGFR1 KD (Figure S1A) (23,24). The lack of kinase activity of VEGFR1 was attributed to an inhibitory sequence in the JM segment (25) and Asn1050 in the A-loop (26), the molecular mechanism of which is unknown.

**Figure 1.**
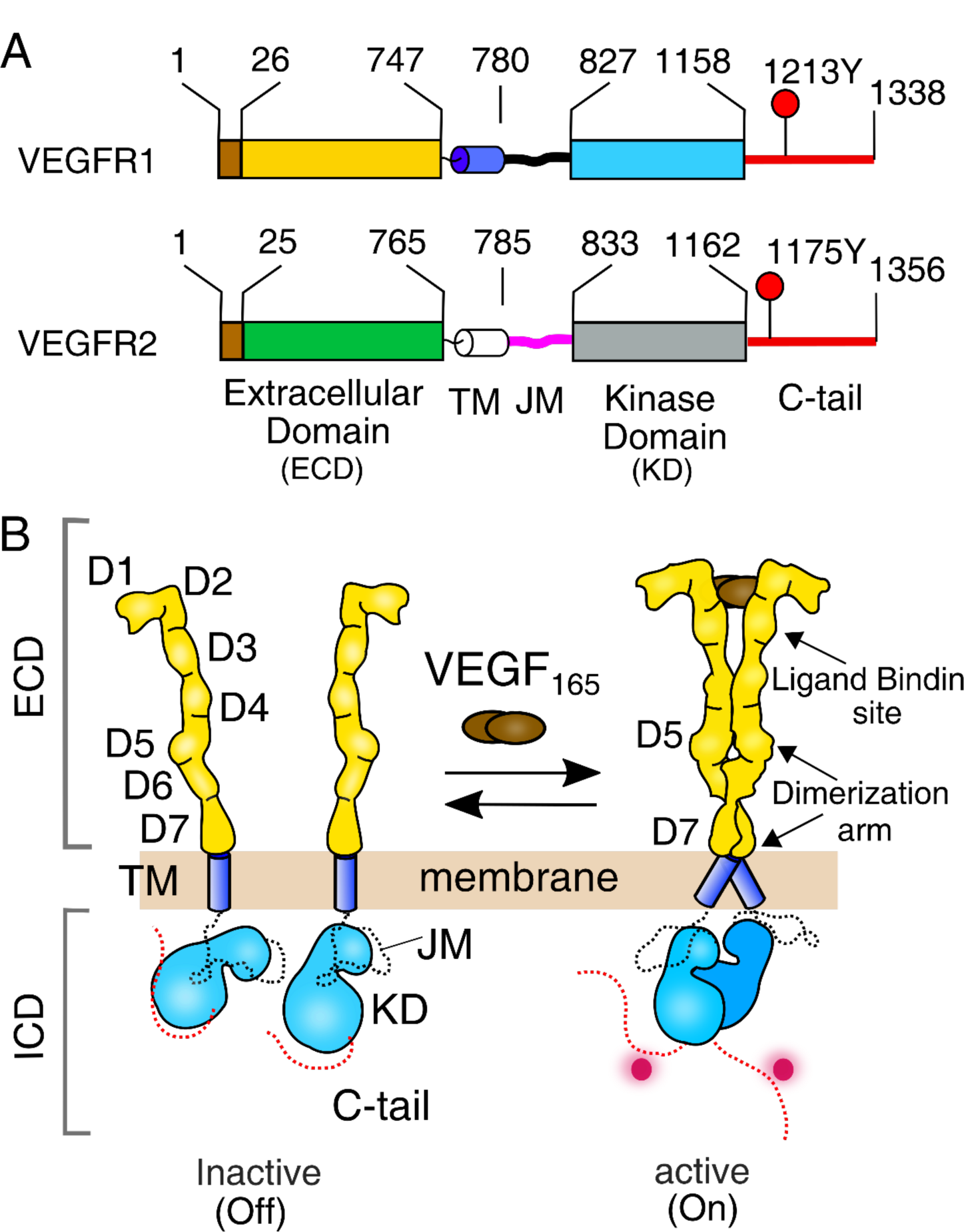
Activation model of VEGFR. **(A)** Schematic representation of domain architecture of VEGFR1 and VEGFR2. The transmembrane and juxtamembrane segment is labelled as TM and JM, respectively. The C-terminal phosphotyrosine residues used for probing kinase activation are labelled. **(B)** Classical model of VEGFR activation in the presence of ligand (VEGF_165_). (Sarabipour et al., eLife,2016) See Figure S1

In contrast, the indispensability of VEGFR1-signaling in the regulation of hematopoietic cell function and the development of pathophysiological conditions is puzzling (5,7). Ligand-dependent activation and VEGFR1-mediated cell-signaling regulate diverse physiological functions(27–30). Overexpression or deregulation of VEGFR1 is linked to several cancers and cancer-associated pain, retinopathy, tumor survival, and autoimmune disorders (31–37). The mechanism of how VEGFR1 autoinhibition is released under pathological conditions is an open question.

To gain further insight, we investigated the ligand-independent and ligand-dependent activation of VEGFR1 and VEGFR2 on the plasma membrane by a single-cell assay using fluorescence microscopy. Our data revealed that, unlike VEGFR2, VEGFR1 does not show concentration-dependent autophosphorylation in the basal state (without a ligand) and is transiently phosphorylated upon ligand stimulation. We decipher that an electrostatic latch in the JM-S and an H-bond between a tyrosine residue in the JM-B and C-helix in VEGFR1 together constitute a JM inhibition that likely stabilizes the inactive JM-in conformation. Slow release of the JM inhibition makes the VEGFR1 autophosphorylation inefficient. Finally, we proposed a mechanism explaining how a delicate balance between kinase and protein tyrosine phosphatase (PTP) maintains the VEGFR1 signaling constitutively off in the basal state.

## Results and Discussion

### VEGFR1 does not show concentration-dependent activation without ligands at the plasma membrane

Ligand-independent activation of RTKs is a key signature of several forms of cancer and manifestation of drug resistance (38–44). Receptor density at the plasma membrane is an important determinant of ligand-independent activation of RTKs (45–50). The density-dependent activation of RTK was explained by an equilibrium shift model between multiple receptor species (51,52). Recent studies showed that VEGFR2 forms a ligand-independent dimer at a physiological concentration on the membrane and is able to autophosphorylate (22). We ask, in the basal state, if VEGFR1 autophosphorylates spontaneously on the plasma membrane.

We begin with a single-cell assay to comparatively study the concentration-dependent activation of VEGFR1 and VEGFR2 at the plasma membrane with and without ligand stimulation, respectively (Figures 1, S1, and S2). We transiently transfected CHO cell lines with VEGFR1-mCherry or VEGFR2-mCherry constructs and stimulated them with VEGF_165_ (Figure S1 B-C). The transient transfection generates a heterogeneous population of cells expressing a diverse concentration of receptors on the plasma membrane. Since the localization of VEGFR family kinases does not solely restrict to the plasma membrane (53,54), we focused on the peripheral regions of the cell for our study (Figure S1G). The activation of the receptor at the membrane was probed by determining the phosphorylation level of Y1213 or Y1175 for VEGFR1 or VEGFR2, respectively, with specific antibodies (55–57). We observed that the unligated VEGFR2 did not autophosphorylate Y1175 at low receptor concentrations but phosphorylates spontaneously at higher receptor concentrations (52) (Figures 2A, B, and E). VEGFR2 linearly phosphorylates Y1175 upon ligand stimulation, suggesting the phosphorylation is independent of receptor concentration at the plasma membrane.

**Figure 2.**
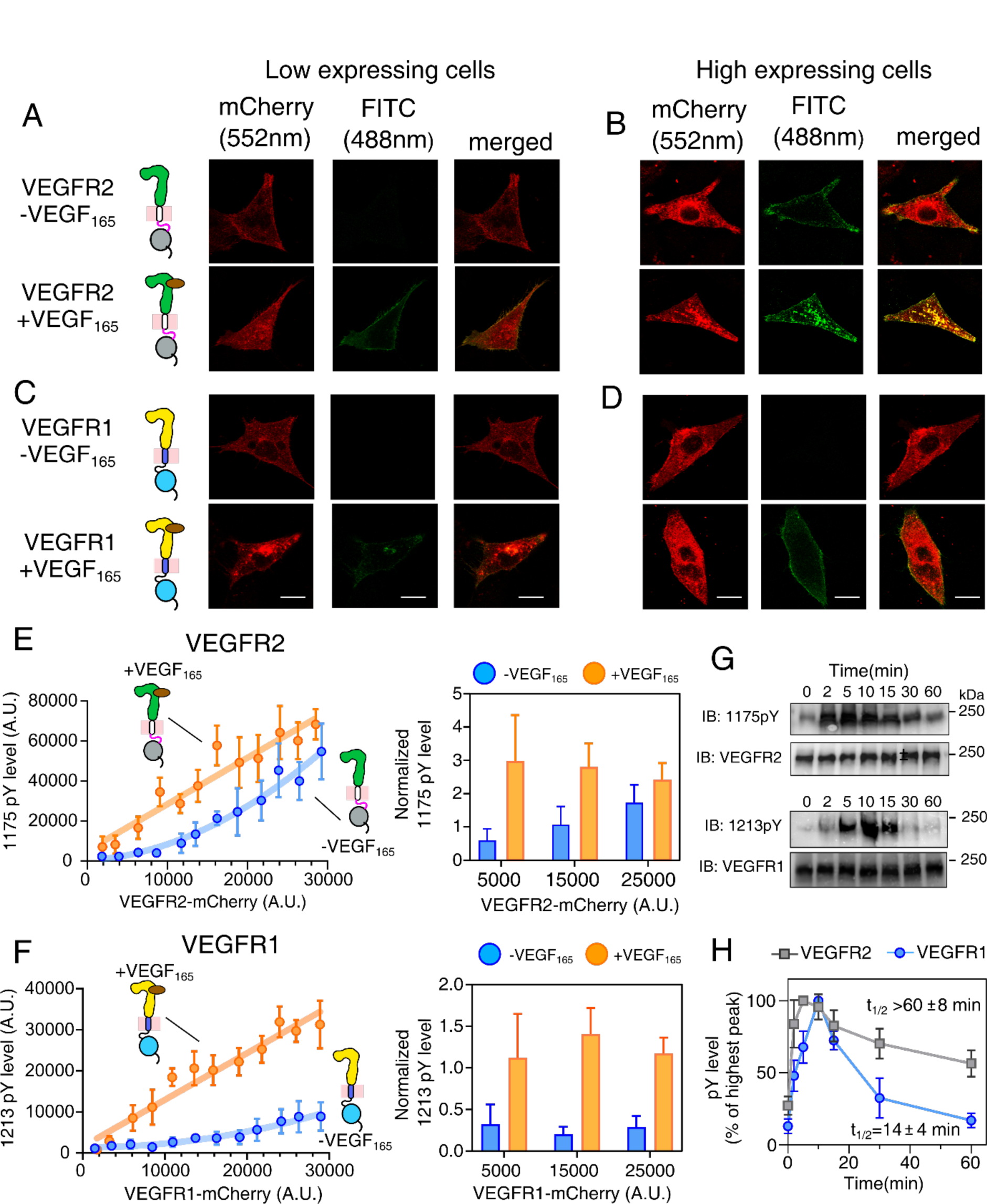
Measurement of ligand-independent and dependent VEGFR activation on the plasma membrane. **(A-D**) Confocal images of VEGFR2 or VEGFR1 fused to mCherry in a low (panel A and C) and high (panel B and D) expressing CHO cell lines. The VEGFR expression level is shown in red (α_ex_= 552nm, α_em_= 586-651nm), and the phosphorylation status is shown in green (α_ex_= 488, α_em_= 505-531). Scale bar = 10 μm) **(E-F**) The expression level of VEGFR2 (panel E) or VEGFR1 (panel F) is plotted against the phosphorylation level of the corresponding tyrosine residues at the C-terminal tail. Individual data points in the left panel represent the mean expression and phosphorylation level for the binned cells. The orange line represents the linear fitting of the individual data points in the ligand-dependent activation. The blue line in panel E represents the second-order polynomial fitting of the individual data points in the ligand-independent activation. In panel F, the blue line is the guiding line. The right panel represents the bar plots of respective phosphotyrosine levels normalized against the VEGFR expression levels in the indicated range. The error bar shows the standard deviation of data points. **(G)** The immunoblot shows the representative phosphorylation level of VEGFR1 or VEGFR2 at the indicated time points after activating the transfected CHO cell line with 50nM VEGF_165_. **(H)** The plot of the phosphorylation level of respective C-terminal tyrosine residue as a function of time. The phosphorylation level is analyzed from the densitometric measurement of the Western blot shown in panel G. The t_1/2_ is determined by fitting the decay of the highest intensity observed to exponential decay. The error bar shows the standard deviation from three independent experiments. See Figures S1 and S2

In the single-cell assay, the VEGFR1 also linearly phosphorylates Y1213 when stimulated with VEGF_165_ (Figures 2D and F). However, the fraction of tyrosine phosphorylated by VEGFR1 is significantly lower than VEGFR2 (Figure S2F). Unexpectedly, we observed that VEGFR1 did not show any ligand-independent autophosphorylation of Y1213, even at the highest receptor concentration measured in our studies (Figures 2C and F). To rule out if the lack of ligand-independent activation of VEGFR1 is not cell-dependent, we repeat the assay by transiently transfecting VEGFR1 to COS-7 and a macrophage cell line (RAW264.7) (Figures S1D-F, S2A-E). We observed a similar phosphorylation profile of Y1213, as seen in the CHO cell line. Suggesting that the lack of ligand-independent activation of VEGFR1 is an intrinsic property of the receptor and not an artifact. We then ask: why does VEGFR1 phosphorylate a lower fraction of tyrosine residues than VEGFR2 upon ligand stimulation? What is the molecular basis that constitutively inactive VEGFR1 in the basal state?

### Phosphorylation of VEGFR1 is transiently stable compared to VEGFR2

To understand why VEGFR1 and VEGFR2 are phosphorylated differentially upon ligand stimulation, we next studied the phosphorylation kinetics and half-life of phosphotyrosine residue (Figures 2G-H and S2 G-I). We determined the phosphorylation level of Y1213 or Y1175 in VEGFR1 or VEGFR2, respectively, by immunoblotting over a period of time after ligand stimulation. The phosphorylation kinetics (Figure 2H, S2H, and Table S4) shows that VEGFR1 is phosphorylated slower (rate = 0.07± 0.01 A.U/min) than VEGFR2 (rate = 0.17± 0.02 A.U/min). We also note that the phosphotyrosine (Y1213) in VEGFR1 is transiently stable (t_1/2_ = 14 ± 4 min) compared to sustained phosphorylation of Y1175 in VEGFR2 (t_1/2_ > 60 ± 8 min) (Figure 2H, S2I and Table S4). We speculate that the slow phosphorylation rate and transient stability of phosphotyrosine residue in VEGFR1 may contribute to a lower fraction of phosphorylated tyrosine residue (Figure S2F). We ask why the VEGFR1 phosphorylation is transient.

The ligand bias by the ECD dimer is known to generate differential signaling output in RTK (58,59) and is a crucial determinant for deciding the cell fate (60). A transient versus sustained Erk activation generated by stimulating EGFR with two different ligands, EGF or NGF, switches the signaling outcomes to differentiation from the proliferation (61,62). Recent studies suggest that the subtle difference in the ligand-induced ECD dimer may determine the differential signaling output in EGFR (58,59). In contrast, VEGF-A induces similar conformation for the ECD of VEGFR1 and VEGFR2, respectively (Figure S3A) (14,20,21). We ask how a common ligand induces different outputs in two homologous VEGF receptors is unknown.

### Deletion of ECD does not constitutively activates the VEGFR1

The ligand-independent activation of VEGFR is obstructed by electrostatic repulsion between the Ig-like domain (D4-7) in the ECD dimer interface (Figure S3 A) (63–65). Despite that, the C482R mutation in the D5 of VEGFR2, linked to infantile hemangioma(66), constitutively activates the kinase by stabilizing a ligand-independent dimer (22). A similar pathogenic cysteine to arginine substitution was reported for FGF receptors (49,67). This suggests that a conserved ligand-independent activation mechanism prevails in RTKs carrying similar Ig-like ECD fold. However, no such mutation has been reported for VEGFR1. We next investigated if mutating the homologous C482 to arginine constitutively activates VEGFR1. We replace C471R in the D5 of VEGFR1 and determine its activation (Figure 3A-B). As expected, the VEGFR2 C482R mutant is constitutively activated and linearly phosphorylates Y1175 even without a ligand (Figure 3C and S3C, E). Surprisingly, the VEGFR1 C471R mutant, in the unligated state, is constitutively autoinhibited (Figure 3D and S3B, D). We wonder if the inability to dimerize renders the VEGFR1 C471R mutant inactive.

**Figure 3.**
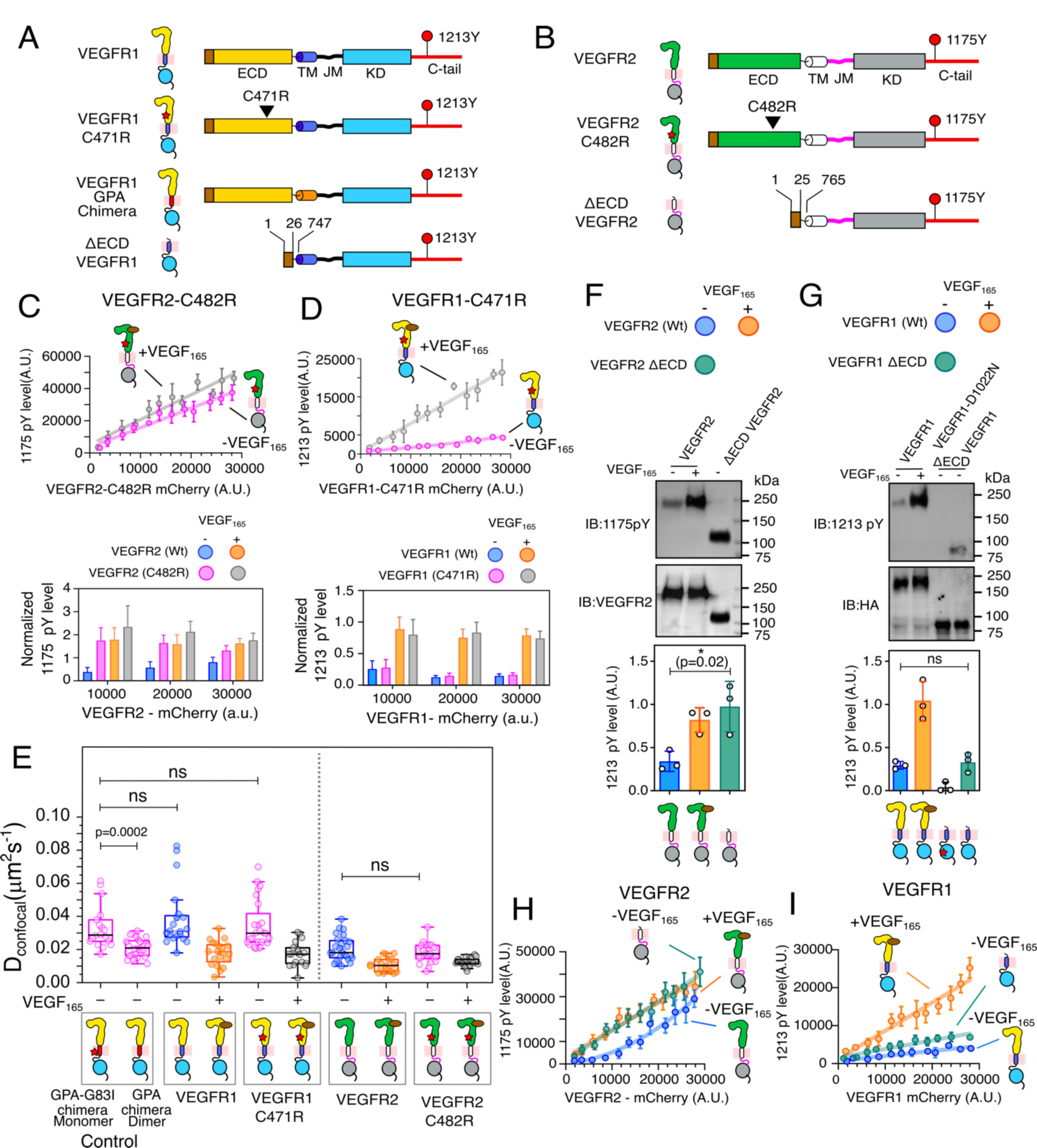
Probing the role of ECD in stabilizing the VEGFR1 autoinhibited state. **(A-B**) Schematic representation of VEGFR1 (panel A) and VEGFR2 (panel B) constructs used in this study. **(C)** The top panel plots the Y1175 phosphorylation level against the expression level of the constitutively activated C482R mutant of VEGFR2 in the presence or absence of VEGF_165_. In the bottom panel, the normalized Y1175 phosphorylation level at the indicated receptor expression level in the C482R mutant and wild-type (wt) VEGFR2 is plotted as a bar graph. **(D)** The upper panel plots the Y1213 phosphorylation and VEGFR1-C471R expression in the presence and absence of ligand. The bottom panel shows the bar plots of the normalized Y1213 phosphorylation level of VEGFR1-C471R and the wt, respectively, at the indicated receptor expression level. **(E)** The diffusion coefficient measured from FRAP studies of indicated constructs of VEGFR1 and VEGFR2 in the presence and absence of VEGF_165_ are plotted. VEGFR1-GPA chimera and VEGFR1-GPA-G83I chimera represent dimer and monomer controls, respectively. Each data point in the box plot reflects the diffusion coefficient of the selected cell, and the black line indicates the mean value. (For each construct, n=25-30 cells) **(F-G**) The immunoblots show the effect of ECD deletion (Δ ECD) on the phosphorylation of Y1175 in VEGFR2 (panel F) and Y1213 in VEGFR1 (panel G). The densitometric analysis of the Western blot is shown below in each panel. The phosphorylation level for each construct is normalized against the respective receptor expression level. In panel G, VEGR1-D1022N represents a kinase-dead mutant. The concentration of the VEGFR1 is determined using an antiHA antibody. (**H-I**) The plot of the phosphorylation level of Y1175 in VEGFR2 (panel H) and Y1213 in VEGFR1 (panel I) against the indicated receptor expression level in the presence and absence of the ligand. See Figures S3 and S4

We next study the oligomeric states of VEGFR1 and VEGFR2 by fluorescence recovery after photobleaching (FRAP) experiment (Figure 3 and S4) (68). The oligomeric status of VEGFR constructs was determined from the diffusion coefficient (*D_confocal_*) derived from the rate of fluorescence recovery at the bleached spot on the plasma membrane. In our experiment, we used two chimeric constructs of VEGFR1 as a monomer (named VEGFR1-GPA-G83I) and dimer (named VEGFR1-GPA) control, where the TM helix is replaced by glycophorin-A (GPA) G83I mutant and wild-type GPA, respectively (Figure 3A) (69,70). The observed increase in the *D_confocal_* for the VEGFR1-GPA-G83I mutant (0.033 ± 0.013 µm^2^s^-1^) confirms a dimer-to-monomer transition on mutating TM segment in the VEGFR1-GPA chimera (0.021 ± 0.005 µm^2^s^-1^) (Figure 3E, S4G, and Table S3). We first turned to VEGFR2 to determine the *D_confocal_* for the wild-type and C482R mutant in the presence and absence of VEGF_165_, respectively (Figure 3E, S4E-F, and Table S3). Overall, *D_confocal_* for VEGFR2 agrees with the recently published data (22). The wild-type VEGFR2 (*D_confocal_* = 0.021 ± 0.008 µm^2^s^-1^) tends to form a ligand-independent dimer, which explains the concentration-dependent activation of VEGFR2 in the basal state (Figure 2E). The ligand binding reorients the ECD and induces dimerization (*D_confocal_* = 0.011 ± 0.004 µm^2^s^-1^) mediated by a homotypic interaction between D4, D5, and D7 (Figure 3E, S3A, and S4C-F) (14,20,21). Our data shows that the VEGFR2 C482R mutant forms a stable ligand-independent dimer (*D_confocal_* = 0.017 ± 0.006 µm^2^s^-1^) that spontaneously activates the KD (Figure 3C, E and Table S3)(22).

We then evaluated the dimerization propensity for the VEGFR1 constructs and made the following observations (Figure 3E, S4, S5 A-D, and Table S3). 1) The VEGFR1 does not dimerize in the absence of a ligand (*D_confocal_* = 0.038 ± 0.018 µm^2^s^-1^), and the ligand binding induces receptor dimerization (*D_confocal_* = 0.018 ± 0.007 µm^2^s^-1^) (Figure 3E, and S4A-B, E-F). 2) Formation of the ligand-dependent dimer is independent of kinase activity as suggested by the *D_confocal_* (0.016 ± 0.006 µm^2^.s^-1^) of the kinase-dead mutant (D1022N) (Figure 5A-B). 3) The C471R mutation does not induce spontaneous dimerization (*D_confocal_* = 0.035 ± 0.014 µm^2^s^-1^). The mutant only dimerizes upon VEGF_165_ binding (*D_confocal_* = 0.017 ± 0.007 µm^2^s^-1^). In summary, our data indicate that VEGFR1 remains predominantly an inactive monomer in the basal state, and the ECD is likely the dominant negative regulator of VEGFR1 activation. Therefore, removing ECD inhibition might spontaneously activate the VEGFR1, as observed in many RTKs (71–74) and is often linked to pathological manifestations (75).

To test this, we measured the autophosphorylation of Y1213 and Y1175 in the ECD-deleted (ΔECD) construct of VEGFR1 and VEGFR2, respectively (Figure 3F-I). As shown previously (74), the VEGFR2 ΔECD construct was constitutively activated (Figure 3F) and linearly phosphorylates Y1175 in the single-cell assay (Figure 3H). Counter-intuitively, the deletion of ECD did not activate the VEGFR1 even at the higher receptor concentration (Figure 3G, I). We speculate that the TM-JM segment connecting the ECD and KD (Figure 1) may be constitutively inhibiting the basal activation of VEGFR1.

### The transmembrane domain does not drive ligand-independent VEGFR1 activation

The TM segment is a major driving force for RTK dimerization. The dynamic equilibrium between receptor dimer and monomer is rotationally coupled to the orientation of the TM segment (76,77). VEGFR2 TM segment adopts two dimer structures, ligand-independent and ligand-dependent (Figure 4D) (22,78). The sequence comparison between the VEGFR1 and VEGFR2 shows that the residues at the ligand-independent dimer interface are conserved (Figure 4D). In contrast, the residues at the ligand-dependent dimer interface are not conserved. We speculate that T763 and C764 make the VEGFR1 TM segment incompatible with a ligand-independent dimer. Without a ligand, the receptor remains in the monomeric state. Ligand binding favors the TM structure towards a ligand-dependent dimer. To test that, we mutated the T761, T763, or C764 individually in VEGFR1 to the corresponding residue in VEGFR2 and measured the Y1213 phosphorylation (Figure S5E). We also replaced the TM segment of VEGFR1 with VEGFR2 in the full-length and ΔECD construct of VEGFR1 (Figure 4A-C, E, and S5E). We observed that none of the mutant and TM chimeric constructs could activate the VEGFR1 ligand independently. The FRAP analysis suggests that the full-length VEGFR1-TM^VEGFR2^ chimera has a higher propensity to form a ligand-independent dimer (*D_confocal_* = 0.022 ± 0.007 µm^2^s^-1^) than the wild-type VEGFR1 (Figure 4F, S5G-H, and Table S3). Suggesting the TM segment of VEGFR1 is a weak dimerization motif compared to VEGFR2 in a ligand-free state. To determine if a stronger TM dimerization motif could spontaneously phosphorylate Y1213 independent of VEGF_165_ stimulation, we turned to the VEGFR1-TM^GPA^ chimera (Figure S4I). We observed that even the VEGFR1-TM^GPA^ chimera could not phosphorylate the Y1213 constitutively. The inability of the VEGFR1-TM^VEGFR2^ or VEGFR1-TM^GPA^ dimer to activate the KD ligand independently is counterintuitive. These data indicate that the regulatory elements downstream of the TM segment may constitutively autoinhibit VEGFR1 in the basal state. Therefore, we replaced the JM or TM-JM segment of VEGFR1 with the VEGFR2 in the ΔECD background. Replacing the JM segment spontaneously activates the kinase and linearly phosphorylates Y1213 (Figure 4B-C and S5F). Together we conclude that the JM segment is a key regulator of VEGFR1 activation.

**Figure 4.**
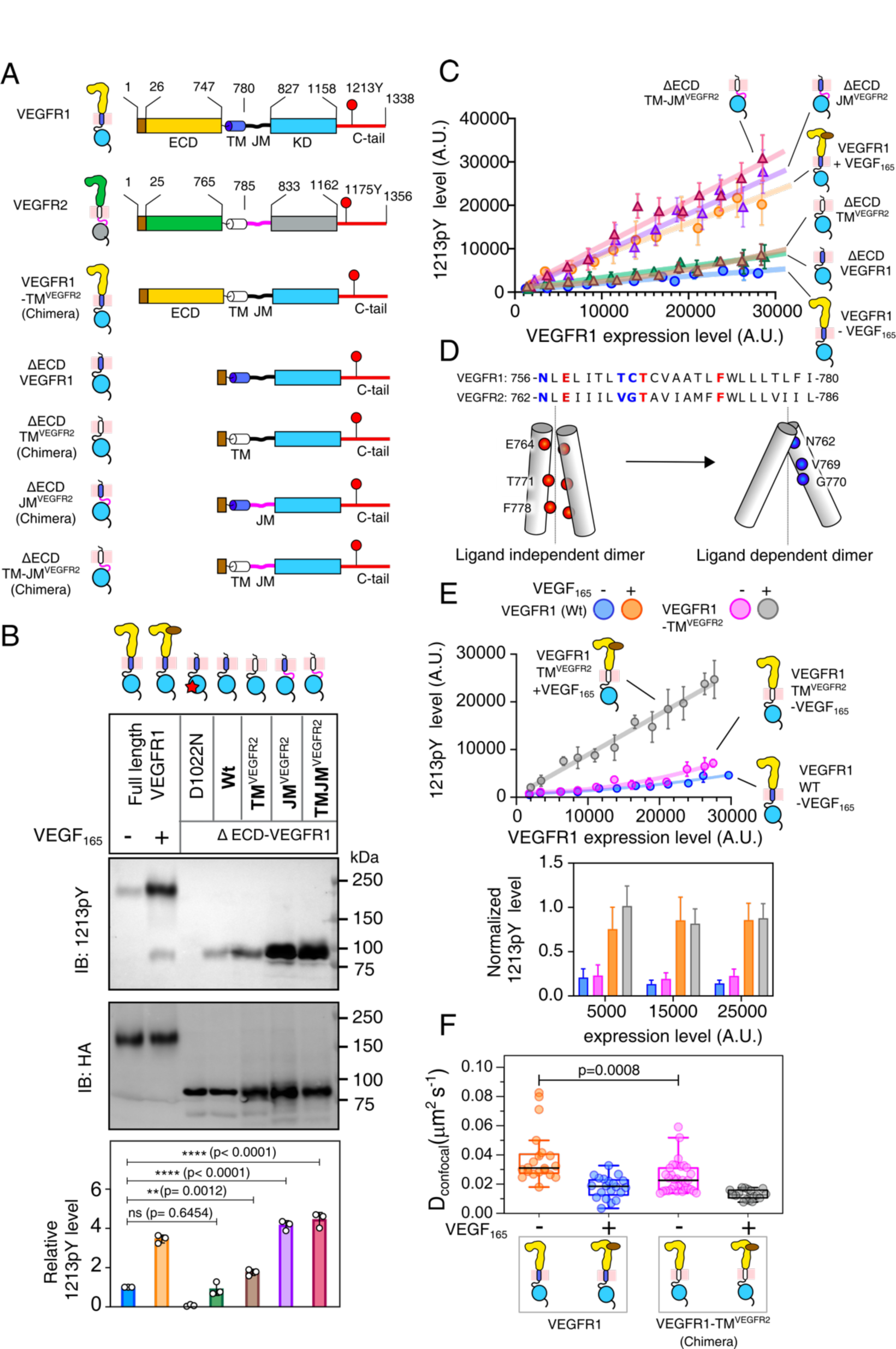
Functional analysis of TM and JM segments in ligand-independent activation of VEGFR1. **(A)** Schematic representation of VEGFR1 and VEGFR2 constructs used in this study **(B)** Immunoblot showing the phosphorylation of Y1213 in the indicated constructs of VEGFR1. The expression level of the VEGFR1 is determined using an antiHA antibody. The bar plot in the lower panel represents the relative Y1213 phosphorylation level determined from densitometric analysis. The data represent the mean ± SD (n=3). **(C)** The plot of the Y1213 phosphorylation level against the expression level of VEGFR1 ΔECD and wt from the single-cell assay. **(D)** Sequence alignment of TM segment of VEGFR1 and VEGFR2. The amino acid residues at the VEGFR2 ligand-independent and dependent dimer interface are colored red and blue, respectively. Below, is the cartoon of the ligand-independent and dependent VEGFR2 TM dimer (Sarabipour et al., eLife,2016; Manni et al., Structure,2013). **(E)** The upper panel plots Y1213 phosphorylation and the expression level of indicated VEGFR1 constructs. The bottom panel shows the bar plot of the normalized Y1213 phosphorylation level for the indicated VEGFR1 chimera and the wt, in the presence and absence of ligand. **(F)** The dimerization propensity of indicated VEGFR1 constructs is probed from the diffusion coefficient measured by the FRAP experiment. (For each construct, n=20-30 cells). See Figures S5

### The electrostatic latch stabilizes the inactive conformation of the JM segment

A repressor sequence present in the JM segment of VEGFR1 is known to inhibit the downstream signaling and cell migration constitutively (25). The JM segment of VEGFR1 and VEGFR2 are homologous and have minor differences in the amino acid sequence (Figure 5A). In the autoinhibited state, the KD of VEGFR1 (PDB ID: 3HNG) and VEGFR2 (PDB ID: 4AGC) adopts a JM-in-like inactive conformation, found in the PDGFR family of kinases (Figure S6A) (16–18). The JM-B segment is buried deep into the catalytic site, stabilizing the folded conformation of the activation loop and preventing the rearrangement of the N-and C-lobe to an active state. The conformation of the JM-Z region sets the direction of the rest of the JM segment inward to the KD. In spite of sharing a high degree of sequence and structural homology, it is not clear how the JM segment of VEGFR1 is differentially regulated from VEGFR2.

**Figure 5.**
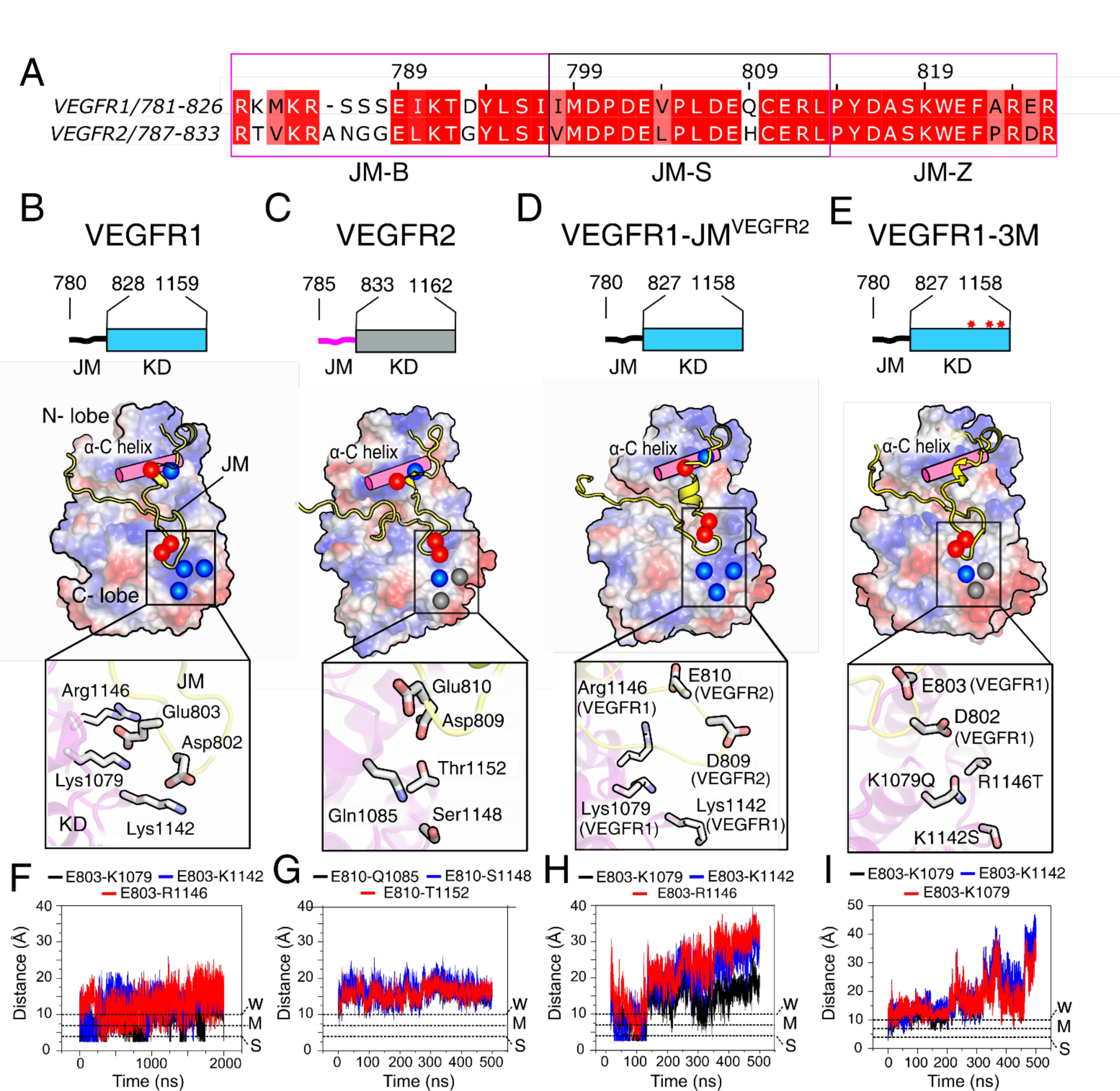
Structural analysis of JM inhibition by MD simulation. **(A)** Sequence alignment of VEGFR1 and VEGFR2 JM segments. **(B-E)** The upper panel shows the schematic diagram of the JM-KD construct used for MD simulation. The bottom panel is the space-filled model of VEGFR1 and VEGFR2 constructs. The electrostatic surface potentials are colored blue and red for the positive and negatively charged sidechains, respectively. The polar uncharged residues are colored grey. The electrostatic interactions (electrostatic latch) between JM and the C-lobe of the KD are shown in the inset. **(F-I)** The pairwise interatomic distances determined for the residues in the electrostatic latch are plotted against time for the constructs described in panels B-E, respectively. The horizontal lines represent the distance cut-off for the weak, medium, and strong electrostatic interactions (89). See Figures S6

To find answers, we revisited the structure of the VEGFR and PDGFR family of kinases. In the crystal structure of VEGFR, the JM-S segment was unresolved. We model the JM-S segment of VEGFR1 and VEGFR2 based on the inactive structure of PDGFR (PDB ID: 5K5X) (Figure S6A). The structural evaluation revealed two key aspects: First, the JM-S segment carries an overall negative charge. Second, the C-lobe of VEGFR1 has a positive charge patch, which is absent in the other VEGFR and PDGFR family (Figure 5A-C). In our model, the positive charge residues (K1142, K1079, or R1146) in the C-lobe of VEGFR1 KD form salt bridges with the negatively charged residues (D802 and E803) in the JM-S segment (Figure 5B). In VEGFR2, the corresponding residues in the C-lobe do not form salt bridges with the JM-S segment (Figure 5C and S6J). We hypothesized that the unique salt bridge between the JM-S and the C-lobe acts as an electrostatic latch that stabilizes the JM-in conformation of VEGFR1, rendering it constitutively inactive in the basal state.

To test our hypothesis, we use molecular dynamics simulation to find the relative stability of the electrostatic latch in VEGFR1, and VEGFR2, respectively (Figure 5F-G, S6B-C, J-K). Our analysis of the distance between the ion pairs suggests that the electrostatic latch in VEGFR1 is considerably more stable than VEGFR2 (Figure 5F-G and S6J-K). We observed that the E803 and D802 in the JM-S segment of VEGFR1 maintained electrostatic contact with the respective C-lobe residues during the simulation. We conclude that the electrostatic latch is an integral component of the autoinhibited VEGFR1 structure and may regulate the transition between an inactive to an active conformation.

### Removing JM inhibition increases the basal activation of VEGFR1

To evaluate the structure and function of the electrostatic latch, we interrogate two VEGFR1 constructs, VEGFR1-JM^VEGFR2^ chimera, and triple mutant (3M) (where positively charged residues in the C-lobe K1142S, K1079Q, and R1146T are mutated to the corresponding residues in VEGFR2) (Figure 5D-E, S6D-E, and 6A). Our structural model shows that the electrostatic latch is broken in the VEGFR1-JM^VEGFR2^ and 3M construct (Figure 5 D-E, H-I, and S6 D-E). We speculate that perturbing the electrostatic latch may destabilize the autoinhibitory interaction of the JM-B. Thus, replacing the VEGFR1 JM segment with VEGFR2 or the triple mutant (3M) may restore ligand-independent activation of VEGFR1. In the single-cell assay, replacing VEGFR1 JM or TM-JM segments with VEGFR2 (Figure 6B-C and S7C-D) restores the concentration-dependent autophosphorylation in the basal state. The 3M mutant partially restored the Y1213 phosphorylation, suggesting a critical role for the electrostatic latch in stabilizing the inactive JM-in structure (Figure 6D). However, the complete restoration of the ligand-independent VEGFR1 autophosphorylation might require additional JM restraint to be removed.

**Figure 6.**
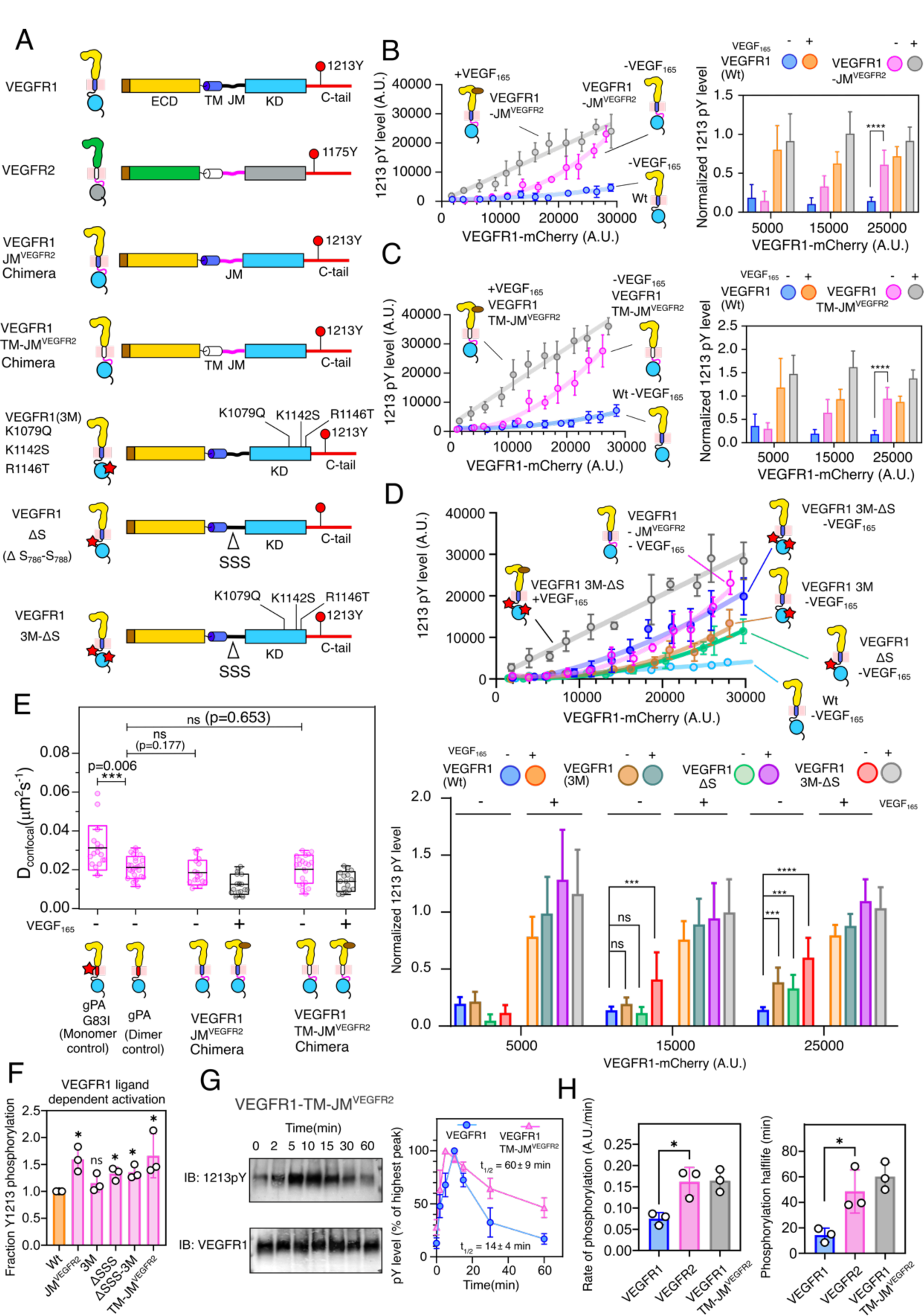
Functional study of JM segment in regulating concentration-dependent activation of VEGFR1. **(A)** Schematic representation of chimeric constructs and mutants of VEGFR1 used in this study (**B, C**) Concentration-dependent activation of VEGFR1 constructs is determined using a single-cell assay in the presence and absence of ligands. The bar plots on the right represent the normalized phosphorylation levels of different chimeric constructs with their respective receptor expression level. **(D)** The upper panel plots Y1213 phosphorylation and the expression level of indicated VEGFR1 constructs. The bottom panel shows the bar plot of the normalized Y1213 phosphorylation level for the indicated VEGFR1 chimera and the wt, in the presence and absence of ligand. **(E)** The dimerization propensity of indicated VEGFR1 construct is probed from the diffusion coefficient measured using the FRAP experiment. (For each construct, n=25-30 cells). **(F)** The relative fraction of phosphorylated Y1213 for the indicated VEGFR1 construct upon VEGF_165_ stimulation is shown as a bar diagram. The fraction phosphorylated was obtained from the slope of the ligand-dependent activation of the respective VEGFR1 construct, as described in panels B, C, and D, and normalized against the wt data. **(G)** The left panel is a representative immunoblot of ligand-dependent Y1213 phosphorylation of VEGFR1-TM-JM^VEGFR2^ chimera measured over indicated time points. The right panel shows the densitometric analysis of the Y1213 phosphorylation level at the indicated time points for the wt (blue) and chimeric construct (magenta) of VEGFR1. The phosphorylation level at each time point is normalized against the highest intensity observed for the respective data set. **(H)** The rate of phosphorylation (left panel) and phosphorylation t_1/2_ (right panel) of the indicated VEGFR constructs are determined from the densitometric analysis of ligand-dependent activation, as described in Figure 2G-H and Figure 6D. See Figures S2 and S7.

To investigate why perturbing the electrostatic latch does not fully restore the ligand-independent activation of VEGFR1, we revisited the JM-in structure of VEGFR2. We observed that the conserved Y801 in the JM-B region of VEGFR2, which forms an H-bond with the critical glutamic acid residue in C-helix in the PDGFR (16,79), is moved out of the catalytic site and does not interact with the C-helix (Figure S7A-B). The JM-B segment of VEGFR1, which is shorter by one residue than VEGFR2, is unresolved in the crystal structure (PDB ID 3HNG) (Figure 5A). Based on structural modeling, we predict that moving the corresponding Y794 in VEGFR1 to (-)1 position may place the Y794 in the catalytic site allowing it to interact with the glutamic acid residue in C-helix (Figure S7A-B). We hypothesize that if the Y794 is moved out of the catalytic site and simultaneously removing the electrostatic latch may activate the VEGFR1 ligand independently. Using the single-cell assay, we determined the ligand-independent activation of VEGFR1 in a ΔS mutant (where three consecutive serine residues, 786-788 at the JM-B, are removed) and a ΔS mutant in the 3M background (Figure 6A). We observed that the ΔS mutant could partially restore the Y1213 phosphorylation without ligand (Figure 6D). To our delight, the ΔS mutant introduced in the 3M background restored the ligand-independent VEGFR1 activation to a level comparable to the VEGFR1-JM^VEGFR2^ chimera (Figures 6D and F). We ask if removing the JM inhibition is enough to remodel the phosphorylation kinetics of VEGFR1.

### Removing JM inhibition remodels transient phosphorylation of VEGFR1 to sustained **phosphorylation**

The FRAP experiment shows an increased dimeric propensity for the VEGFR1-JM^VEGFR2^ (*D_confocal_*

= 0.0205 ± 0.0059 µm^2^s^-1^) and VEGFR1-TMJM^VEGFR2^ (*D_confocal_* = 0.0202 ± 0.007 µm^2^s^-1^) chimera (Figure 6E, S7I-J, and Table S3). Suggesting the JM-in conformation of VEGFR1 is incompatible with the ligand-independent TM dimer. Stimulating VEGFR1-JM^VEGFR2^ (*D_confocal_* = 0.0127 ± 0.0047 µm^2^s^-1^) and VEGFR1-TMJM^VEGFR2^ (*D_confocal_* = 0.0131 ± 0.005 µm^2^s^-1^) with VEGF_165_ induces the receptor oligomerization and also increases the relative fraction of Y1213 phosphorylation compared to wild-type VEGFR1 (Figure 6E, F, S7I-J, and Table S3). Removing the JM inhibition in the VEGFR1-TMJM^VEGFR2^ chimera increases the rate of Y1213 phosphorylation (0.16 ± 0.03 A.U/min). It also remodels the phosphorylation half-life from transient to sustained (t_1/2_ = 48 ± 13.8 min) (Figure 6G-H, S7 G-H and Table S4). We conclude that the multiple interactions between the JM segment and the kinase core are the dominant force in autoinhibiting the VEGFR1 constitutively in the basal state. Nevertheless, after an extensive literature review, we could find a pathological mutant constitutively activating VEGFR1. How is basal activation of VEGFR1 regulated in various pathophysiological conditions (33,35,80,81)?

### Cellular phosphatase balance modulates VEGFR-1 basal activation

It is now evident that oxidative stress due to reactive oxygen species (ROS) under various pathophysiological conditions or in experimental setup activates VEGFR and other RTKs (80, 82-85). For example, oxidative stress promotes VEGFR1 overexpression and induces ligand-independent cell migration in renal cell carcinoma and human colorectal cancer cell, respectively (80–82). In diabetic patients, ROS generated in hyperglycemia promotes ligand-independent phosphorylation of VEGFR2 (84). We ask if increasing cellular ROS concentration spontaneously phosphorylates VEGFR1. We treated the COS-7 cells transiently expressing VEGFR1-mCherry with H_2_O_2_ to generate ROS (Figure S8A) and determine ligand-independent phosphorylation of Y1213 (Figure 7A). We observed that wild-type VEGFR1 is autophosphorylated upon H_2_O_2_ treatment, but in the kinase-dead mutant, Y1213 was marginally phosphorylated (Figure 7A and S8A). Suggesting, under oxidative stress, VEGFR1 spontaneously autophosphorylates the Y1213 and does not require help from a second tyrosine kinase. In human colorectal cancer cells and hyperglycemia, it may be noted that the VEGFR1 and VEGFR2 phosphorylation is mediated by Src tyrosine kinase (80,84).

**Figure 7.**
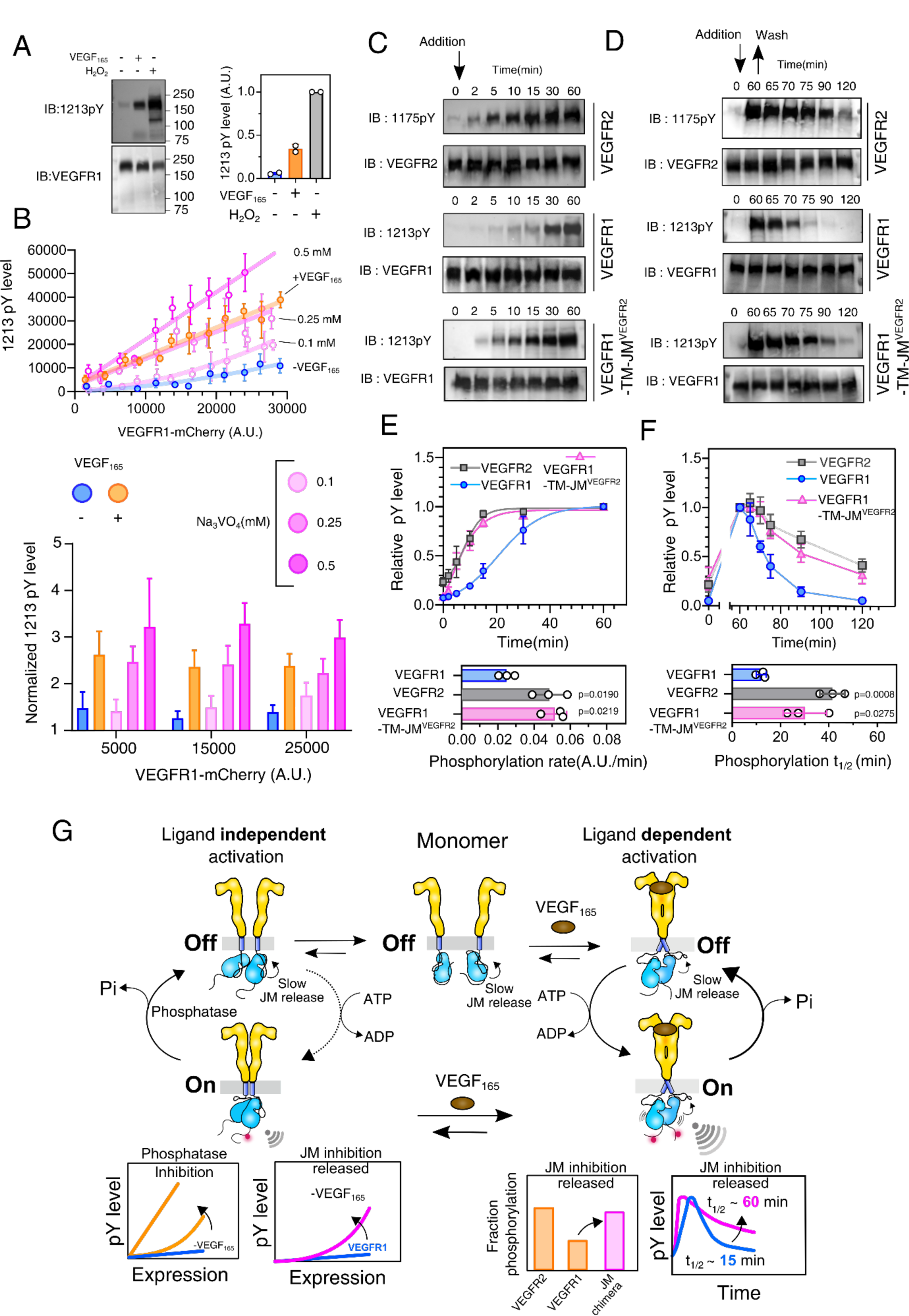
Effect of cellular phosphatase in regulating VEGFR1 ligand-independent autophosphorylation. **(A)** The representative immunoblot (left panel) and densitometric analysis (right panel) of Y1213 phosphorylation by H_2_O_2_. **(B)** Concentration-dependent phosphorylation of Y1213 at the indicated sodium orthovanadate (Na_3_VO_4_) concentration is plotted against the VEGFR1 expression level. **(C)** Representative immunoblot of the tyrosine phosphorylation of VEGFR2 (top), VEGFR1 (middle), and VEGFR1-TM-JM^VEGFR2^ chimera (bottom) measured at the indicated time points upon Na_3_VO_4_ treatment (Shown by down arrow). **(D)** The immunoblot showing the dephosphorylation of indicated phosphotyrosine residue for the VEGFR2 (top), VEGFR1 (middle), and VEGFR1TM-JM^VEGFR2^ chimera (bottom) at the different time points upon Na_3_VO_4_ removal. The up arrow points when Na_3_VO_4_ is washed. **(E and F)** The upper panel is the densitometric analysis of the phosphorylation and dephosphorylation of the respective tyrosine residue in VEGFR constructs described above in panels C and D, respectively. The lower panel is a bar plot of the phosphorylation rate (panel E) and half-life (panel F) of the phosphotyrosine residue in the indicated VEGFR constructs. The error bar shows the standard deviation from three independent experiments. **(G)** The proposed model describes the balance between slow kinase activation due to JM inhibition and phosphatase activity attenuating the VEGFR1 signaling. See Figures S8

The ROS (mainly H_2_O_2_) promotes the phosphorylation of RTKs by inhibiting the PTP (86). We next ask if inhibiting PTP with a nonspecific inhibitor, sodium orthovanadate, would promote the VEGFR1 autophosphorylation. We determined the phosphorylation of Y1213 in transiently transfected CHO cell lines expressing wild-type or kinase-dead mutant (D1022N) of VEGFR1, respectively, after treatment with 1 mM sodium orthovanadate (Figure S8B). Inhibiting PTP significantly increase the phosphorylation of Y1213 compared to ligand stimulation. The kinase-dead mutant shows negligible Y1213 phosphorylation after PTP inhibition. Independently confirms that inhibiting PTP increases the autophosphorylation of tyrosine residues in VEGFR1 and does not require another tyrosine kinase. Further, we observed that VEGFR1 autophosphorylates with increasing orthovanadate concentration and displays concentration-dependent autophosphorylation of Y1213 at a 0.1mM orthovanadate (Figure 7B and S8C). However, in a narrow concentration range of orthovanadate (0.25mM), the Y1213 phosphorylation transit to the linear pattern, as observed on ligand stimulation (Figure 7B). The time-dependent phosphorylation of Y1213 upon orthovanadate treatment shows the wildtype VEGFR1 is a slower kinase (phosphorylation rate = 0.025 ± 0.003 A.U/min) and has a shorter phosphotyrosine half-life (t_1/2_ = 12.0 ± 1.6 min), compared to VEGFR2 (phosphorylation rate = 0.051 ± 0.005 A.U/min, t_1/2_ = 41.4 ± 4.2 min) and VEGFR1-TMJM^VEGFR2^ chimera (phosphorylation rate = 0.048 ± 0.008 A.U/min, t_1/2_ = 30.3 ± 7.3 min) (Figure 7C-F, S8D-E, and Table S4). These data reinforce JM as a master regulator of kinase activation rendering VEGFR1 phosphorylation sensitive to PTP.

### Conclusions

To summarize, we presented a molecular mechanism explaining how VEGFR1 and VEGFR2 are differentially autophosphorylated in the basal state and upon ligand binding (Figure 7G). Unlike VEGFR2, VEGFR1 is constitutively autoinhibited in the basal state, even at high receptor concentrations (Figure 2). Our data support a central role for the JM segment balancing the equilibrium between an inactive and active state of VEGFR1. In the basal state, the electrostatic latch and H-bond interaction steaming from Y794 in JM-B may shift the equilibrium towards the inactive state, thus making VEGFR1 an inefficient kinase. The JM-in conformation of VEGFR1 is incompatible with the ligand-independent TM dimer suppressing spontaneous kinase activation (Figures 5 and 6) (22) Forced dimerization of the TM segment is unable to remove the JM inhibition. We propose that slow kinase activation, and the cellular PTP activity maintains the basal activity of VEGFR1 constitutively inhibited (Figure 7G). The autoinhibition is released either by mutating the JM segment (Figure 6) or inhibiting PTP (Figure 7). We speculate that marginal reduction in PTP activity due to oxidative stress under pathological conditions may be sufficient to stimulate ligand-independent VEGFR1 signaling.

Ligand binding induces receptor dimerization, leading to the rearrangement of the TM-JM segment. The structural rearrangement activates the KD by releasing JM inhibition, causing transient inhibition of PTP due to induction of ROS generation (Figure 7G) (87,88). Slow removal of JM inhibition in VEGFR1 and inactivation of PTP leads to transient phosphorylation of tyrosine residues. Removing the JM inhibition may restore sustained tyrosine phosphorylation in VEGFR1 (Figure 7G). We conclude that the signaling bias of VEGFR1, in basal state or upon ligand stimulation, may be determined by the dynamics of JM inhibition and the phosphatase status of the cell.

## Experimental Procedures

The experimental procedure and material section are provided in the supporting information.

## Supporting Information

The Supporting Information contains Experimental Procedures, Figure S1-S8, and Table S1-S5.

## Acknowledgments

The authors thank Dr. Arnab Gupta and his group members for access to the microscope facility. The authors thank Kaustav Gangopadhyay, Swarnendu Roy, and Subhankar Chowdhury for their help in the initial stage of the project and comments.

## Author Contributions

The manuscript was written through the contribution of all authors. All authors have approved the final version of the manuscript. RD and MPC designed the experiments. MPC, PM, PK, SB, and PKD performed the Biochemical experiments, Imaging, and data analysis. DD did the molecular dynamics simulation and data analysis. RD and MPC wrote the manuscript.

## Funding and additional information

The authors thank research funding from IISER Kolkata, infrastructural facilities supported by IISER Kolkata, and DST-FIST (SR/FST/LS-II/2017/93(c)). This work is supported by a grant from SERB (CRG/2020/000437). Fellowships from CSIR-UGC support MPC and PM; Fellowship from KVPY supported DD.

## Data Availability Statement

All the relevant data are contained within this article and in the supporting information.

