## Supplemental Information for "Molecular basis of VEGFR1 autoinhibition at the plasma membrane"

**Supporting Information**

Corresponding authors

Rahul Das:

**Methods and materials:**

**DNA Constructs**

The cDNA encoding wild-type human VEGFR1 (Residue number 1–1338; pDONR223-FLT1 was a gift from William Hahn & David Root; Addgene plasmid # 23912) was subcloned into pcDNA-mCherry vector between the HindIII and Kpn1 restriction sites. The plasmid encoding human wild-type VEGFR2 tagged with mCherry (cloned in pBE-vector; 108854) was a gift from Kalina Hristova (Johns Hopkins University, Baltimore, MD). All point mutations and deletion constructs were prepared by PCR-based mutagenesis strategy [1]. For the VEGFR1 ECD deletion constructs, Haemagglutinin (HA) tag was incorporated at the 5′ ends. The list of primers used is tabulated in Table S3. In brief, PCR-amplified product at the expected molecular weight was extracted and purified from 0.6% agarose gel and incubated with Dpn1 at 37°C for 3 hrs. The Dpn1 digested PCR product was further purified from the reaction mixture and incubated with Polynucleotide Kinase 4 (PNK4) and Ligase in Tris buffer containing 10 mM $MgCl_2$ and 2 mM ATP. Finally, the ligated mixture was transformed into the *E.coli* Top10 cell. Chimeric constructs were prepared using the Gibson assembly approach [2]. All the chimeric constructs were cloned into BamH1 and Xho1 restriction sites in a pcDNA-mCherry vector (mCherry2-N1 was a gift from Michael Davidson, Addgene Plasmid # 54517)

**Cell culture, Immunofluorescence**

Chinese Hamster Ovary (CHO), African green monkey kidney fibroblast-like cell line (Cos-7), and murine macrophage (RAW264.7) cell lines were cultured in DMEM supplemented with 10% FBS, 50µg/ml penicillin, and streptomycin at 37 °C with 5% $CO_2$. In the single-cell experiments and immunoblotting, VEGFR1 or VEGFR2 was transiently transfected in respective cell lines. In brief, the Cos-7 or the CHO cells were grown on coverslips (10 mm, #1 thickness) up to a confluency of 80% and transfected with Lipofectamine 2000 in serum-free media (Opti-MEM). After 6 hours of incubation, cells were supplemented with complete media containing 10% FBS and allowed to grow for another 12 hours. Alternatively, RAW264.7 cells were nucleofected using the D032 program in the Lonza nucleofector. $4X10^7$ cells were incubated with 4µg of DNA in Amaxa buffer for 5 min at room temperature. Nucleofected cells were resuspended in DMEM supplemented with 15% FBS. Eighteen hours post-transfection, cells were serum-starved by growing in serum-free opti-MEM for another 8 hrs. To activate VEGFR1 or VEGFR2, transfected cells were treated with 100nM $VEGF_{165}$ for 5 mins, then immediately washed with ice-cold PBS two times and fixed by treating with 4% paraformaldehyde (PFA) in PBS for 30 minutes at room temperature. After fixation, cells were washed five times with 1x PBS and permeabilized with 0.2% PBST (1X PBS and 0.2% Triton-X 100)) for 5 min at room temperature. Cells were blocked with 1% BSA for 1 hour, followed by staining with anti-phosphotyrosine antibodies for VEGFR2 or VEGFR1 overnight at $4^0$C. After overnight incubation, coverslips were washed three times with PBST, followed by incubation with a secondary antibody conjugated with FITC for 2 hours at room temperature. Finally, coverslips were mounted with prolonged gold, in between each step.

#### Microscopy and image analysis

Transiently transfected COS7 cells were imaged by an Olympus IX81 epifluorescence microscope. For CHO and RAW264.7 cell line, all images were acquired with the Leica SP8 confocal platform using an oil immersion HC PL APO [3]pinhole was set at 1 AU. Image scanning was done in bidirectional mode at 500 Hz. Expression of mCherry fused VEGFR constructs and the phosphorylation levels was imaged using 40mW/552nm and 20mW/488nm solid-state lasers, respectively. For mCherry, $\lambda_{ex}$ and $\lambda_{em}$ were set to 552nm and 576-651nm, respectively. For FITC, $\lambda_{ex}$ and $\lambda_{em}$ were set to 488nm and 505-531 nm, respectively. Laser power was set at 0.08 mW and 0.04 mW for mCherry and FITC channels, respectively. The background fluorescence was corrected by subtracting the respective fluorescence measured for untransfected cells. Image analysis was done by selecting ROIs drawn manually using ImageJ freehand tool[4], as explained previously [3]. Two sets of ROI were selected; ROI1 was drawn to calculate total cell intensity, and ROI2 was drawn at the peripheral region of the cell to calculate cytoplasmic intensity (Figure S1G). The fluorescence intensity of mCherry or FITC at the membrane ($I_m$) was calculated by:

$I_m = I_{ROI1}-I_{ROI2},$ where $I_{ROI1}$ and $I_{ROI2}$ are fluorescence of mCherry or FITC at the respective ROI.

To analyze the tyrosine phosphorylation as a function of receptor expression level, the intensity of the mCherry channel was binned in the range of 2500 A.U., and the average value of mCherry intensity was plotted against the corresponding average value of FITC intensity. For measuring the normalized phosphorylation level, the intensity from the FITC channel was normalized by the corresponding mCherry intensity for the respective bin, the cells were binned at an intensity range of 10000 A.U.

#### Immunoblotting

The VEGFR kinase domain was activated as described above. After activation, cells were washed with ice-cold PBS before being incubated in RIPA buffer (10 mM Tris-Cl, pH 8.0, 140 mM NaCl, 1 mM EDTA, 0.1% SDS, and 1% Triton X-100) containing protease and phosphatase inhibitors (2 mM Benzamidine, 1 mM PMSF), and then sonicated on ice. Protein samples were prepared by heating them with 5X loading buffer, resolving them on a 6% SDS-PAGE, and blotted onto a PVDF membrane. The blot was blocked with 5% skimmed milk in 1 x TBS and 0.1% TWEEN-20 for 1 hr at RT and then incubated with primary antibody at 4 °C overnight. The unbound primary antibody was removed by washing the blot three times with 1x TBST (0.1% tween-20), followed by incubation with a secondary antibody diluted in 3% skimmed milk. Blot was washed three times with 1x TBST and two times with 1XTBS wash before developing with the Clarity™ Western ECL substrate kit (Bio-Rad). The images were acquired using the Bio-Rad Chemidoc system. The densitometric analysis was performed using ImageJ.

### Cell-based tyrosine phosphorylation kinetic measurements

Chinese Hamster Ovary (CHO) was cultured and transfected with VEGFR constructs as described previously. Before stimulation, cells were serum-starved in Opti-MEM for 8 hours. The VEGFR tyrosine phosphorylation was stimulated by treating with $VEGF_{165}$ or sodium orthovanadate. Samples were collected at the indicated time points and analysed by immunoblotting. To inhibit protein tyrosine phosphatase (PTP), transfected cells were treated with 1 mM sodium orthovanadate and diluted from 100 mM stock solution. The reaction was terminated by lysing the cells at the indicated time points. Densitometry analysis of immunoblots was performed using ImageJ software.

### FRAP Experiment

CHO cells were seeded in a 35 mm glass-bottomed Petri dish at a density of $0.5 \times 10^6$ cells per plate. Transfection was performed as previously described. The cells were grown for eighteen hours post-transfection and were serum starved for six hours before ligand stimulation. Before activating with $VEGF_{165}$, cells were washed with pre-warmed PBS(1x) and activated with 100nM $VEGF_{165}$ in Opti-MEM.

FRAP experiments were conducted using a Leica SP8 confocal microscope at room temperature. Imaging was performed using an HC PL APO CS2 63X/ 1.4 NA objective at 5× digital zoom. The confocal pinhole was set at 1 Airy unit. Image scanning was done in bidirectional mode at 500 Hz. The excitation and emission filters for the mCherry channel were set to 552nm and 576-651nm, respectively. A circular region of radius 1.2 µm (bleached spot) at the cell edge was bleached using a 40mW solid-state laser (100% intensity) for 500 ms, with a pixel dwell time of 1.15 µs. The laser powers for pre-bleach and post-bleach imaging were set at 1% (0.4mW). 10 pre-bleached and 120 post-bleached frames were recorded in 512 X 512 pixel format at 2frame/s.

*FRAP analysis*

The measured raw fluorescence intensity was corrected for background fluorescence and photofading effects. At first, the fluorescence intensity at each frame was measured for the ROI defined by the bleached spot [$F_{ROI}(t)$]. To account for background fluorescence [$F_{bg}(t)$], the fluorescence was measured in a cell-free region of the image in the respective frame. To adjust for observational photofading [$F_{total}(t)$], the fluorescence from the total cell, except the bleached spot, was measured. The raw FRAP data [$F_{ROI}(t)$], were corrected for background and photofading at each frame by:

$$F_{corrected}\ (t) = \frac{F_{ROI}(t) - F_{bg}(t)}{F_{Total}(t) - F_{bg}(t)}$$

Finally, the corrected FRAP ($F_{corrected}\ (t)$) was normalized by corrected prebleach intensity ($F'_{corrected}$):

$$F(t) = \frac{F_{corrected}\ (t)}{F'_{corrected}}$$

$$\left[ F'_{corrected} = \frac{F_{ROI}(t_0) - F_{bg}(t_0)}{F_{Total}(t_0) - F_{bg}(t_0)} \right]$$

(Where $F_{ROI}(t_0)$ , $F_{Total}(t_0)$ , and $F_{bg}(t_0)$ indicate the initial fluorescence intensity from the bleached spot, total cell, and background, respectively)

The effective bleached spot radius ($r_e$) was determined as described previously (Figure S4H) [5]. The half-life of fluorescence recovery ($t_{1/2}$) was determined by fitting corrected fluorescence intensities at indicated time points to the first-order kinetic equation:

$$f(t) = A(1 - e^{-kt}) \ \ \{ \text{where } k = \frac{0.69}{t_{1/2}} \}$$

Finally, the diffusion coefficient ($D_{confocal}$) was derived from the modified Soumpasis equation as described previously[6]:

$$D_{confocal} = 0.25\ \frac{r_e^2}{t_{1/2}}$$

where $r_e$ is the effective bleached spot radius and the coefficient 0.25 was numerically determined [5]

We validated our FRAP experimental setup by comparing the diffusion coefficients ($D_{confocal}$) of ligand-free EGFR-mCherry and VEGFR2-mCherry with previously reported diffusion coefficient measurements using single-molecule tracking. We observed that the diffusion coefficient of EGFR (0.033 $\mu m^2.s^{-1}$) and VEGFR2 (0.020 $\mu m^2.s^{-1}$) (Table S3) determined from Confocal FRAP is in agreement with the average diffusion coefficient value reported for the heterogeneous population of EGFR ( 0.036 $\mu m^2.s^{-1}$ ) [7] and VEGFR2 ( 0.032 $\mu m^2.s^{-1}$) [8], respectively.

**ROS measurement**

ROS measurement was performed using $H_2DCFDA$ as described previously [9]. In brief, Cos-7 cells were seeded into 96 well plates and left to adhere overnight. The ROS was generated by treating the cells with $H_2O_2$. Before treatment, cells were serum-starved for 6 hours and then treated with 1mM $H_2O_2$ diluted in Opti-MEM for 1 hour at 37°C. The cells were washed with pre-warmed PBS(1x) and loaded with 5µM H2DCFDA diluted in Opti-MEM for 30mins at 37°c. The excess $H_2DCFDA$ was removed by washing the cells twice with PBS supplemented with Opti-MEM. Fluorescence intensity was measured using a microtiter plate reader (Biotek synergy h1) at $\lambda_{ex}$ and $\lambda_{em}$ of 485 and 535 nm, respectively.

**Homology Modelling**

The models for the autoinhibited conformation of VEGFR1 (781-1158), VEGFR2 (787-1162), VEGFR1-JM$^{VEGFR2}$ chimera, and VEGFR1 triple mutated (K1079Q, K1142S, and R1146D) were built using iTasser server (https://zhanggroup.org/I-TASSER/)[10]. In all our models, the kinase domain adopts an autoinhibited conformation of C-helix-in and DFG–out. The kinase domain of VEGFR-1 (825-1158) was modeled based on PDB ID 3HNG, and the juxtamembrane segment (781-824) was modeled based on PDGFR structure (PDB ID 5K5X) (Figure S6B). The linker region between the N-lobe and C-lobe (925-991) could not be modeled due to the lack of any homology structure and remained unstructured in the model. The final model, with the highest confidence score as well as a low root-mean-square-deviation (see Table S5, RMSD of 0.589 Å w.r.t 3HNG) with respect to VEGFR1 structure (PDB ID: 3HNG), was selected and the inhibitor N-(4-chlorophenyl)-2-[(pyridin-4-ylmethyl)amino] benzamide was docked at the ATP binding pocket based on 3HNG structure, using PyMOL (DeLano, W. L. ,2009). The structure was further energy minimized and equilibrated before analyzing by MD simulation. A preliminary MD simulation of 100 ns was performed to determine the structural stability of the selected model. The lowest energy structure selected remained stable in the preliminary simulation and was thus used for further studies. The models for the VEGFR1-JM$^{VEGFR2}$ chimera and the triple mutant construct of VEGFR1 were similarly constructed, energy minimized, and evaluated for further structural studies. Similarly, VEGFR2 (PDB ID 4AGC) was used to model the kinase domain (832-1162), and the juxtamembrane segment (787-831) was modeled based on PDGFR structure (PDB ID 5K5X). The model was selected as explained previously, and Axitinib was docked at the ATP binding pocket based on 4AGC structure, using PyMOL.

**Molecular Dynamics simulations**

The MD simulations were run using GROMACS 2019.6[11, 12]. System preparation was done using the CHARMM-GUI server (www.charmm-gui.org/) with TIP3p water molecules[13] and 0.15 M NaCl. The model required 1000 steps approximately for energy minimization. The solvated system was equilibrated for 125 ps at a 1 fs time step, using H-bonds as constraints by implementing Linear Constraint Solver for Molecular Simulations (LINCS) algorithm [14] . At least three independent simulations of 2 μs each for VEGFR1 (wildtype), 500 ns each for VEGFR2 (wildtype), VEGFR1-JM$^{VEGFR2}$ chimera, and VEGFR1 triple mutated were performed using the CHARMM36m force field[15]. To comparatively study the orientation of the juxtamembrane segment with respect to the kinase domain in the crystal structure of cFMS (PDB ID 2OGV) and FLT3 (PDB ID 1RJB), we performed three independent simulation of 200 ns each. The simulations were performed under constant pressure (1 bar) [16] and temperature (300 K) (NPT) [17] and a time step of 2 fs. Potential-shift-Verlet was used for electrostatic and van der Waals interactions with a 12-Å cutoff. The trajectories were visually analysed using VMD[18] , and the structures were visualized in PyMol. RMSDs, RMSFs, and distances were measured using the tools provided in GROMACS.

**Table S1**: List of Antibodies

| Antibodies | Source | |
|---|---|---|
| VEGFR2 Rabbit monoclonal antibody | Cell Signaling Technology, Danvers, MA, USA | Cat # 2479S<br>Lot: 18 |
| Phospho-VEGF Receptor 2 (Tyr1175) Rabbit monoclonal antibody (19A10) | Cell Signaling Technology, (Danvers, MA, USA) | Cat # 2478T<br>Lot:15 |
| VEGFR1 Goat polyclonal antibody | R & D system<br>(Minneapolis, MN, USA) | Cat # AF321<br>Lot: AHT2018011 |
| Phospho-VEGF Receptor 1 (Tyr1213)<br>Rabbit monoclonal antibody | My BioSource<br>(San Diego, CA, USA) | Cat # MBS002880<br>Lot: 07/2021 |
| HA mouse monoclonal antibody | Biolegend<br>(San Diego, CA, USA) | Cat # 901501<br>Lot: B318172 |
| GAPDH rabbit polyclonal antibody | BioBharati Life Science Pvt Ltd | Cat # BB-AB0060<br>Lot: 011501 |
| Mouse HRP Secondary antibody | Cell Signaling Technology, Danvers, MA, USA | Cat # 7076S<br>Lot: 33 |
| Rabbit HRP Secondary antibody | Abcam<br>(Waltham, MA 02453, USA) | Cat# 50095<br>Lot: 2960660 |
| Goat HRP Secondary antibody | Abcam<br>(Waltham, MA 02453, USA) | Cat #Ab6717<br>Lot: GR267728-27 |
| Rabbit FITC conjugated secondary antibody | Abcam<br>(Waltham, MA 02453, USA) | Cat # Ab6885<br>Lot: GR3391568-I |

**Table S2**: List of constructs and primers:

| Construct | Primer | Sequence |
|---|---|---|
| VEGFR2 -C482R | Fwd | GAC AAA CCC ATA CCC TTG TGA AGA ATG GAG AAG TG |
| | Rev | CACTTCTCCATTCTTCACAAGGGTATGGGTTTGTC |
| VEGFR1 -C472R | Fwd | GTT CTG GCA CCC CCG TAA CCA TAA TCA TTCC |
| | Rev | GGAATGATTATGGTTACGGGGGTGCCAGAAC |
| VEGFR1-TM$^{VEGFR2}$ | TM_FP1 | TACCGGACTCAGATCTCGAGATGGTCAGCTACTGGGACACCGG |
| | TM_RP1 | AGAATAATGATTTCCAAGTTAGACTTGTCCGAGGTTCCTTGAACAG |
| | TM_FP2 | AACTTGGAAATCATTATTCTAGTAGGCACGGC |
| | TM_RP2 | AGACCTTTTCATTTTTCGTAGGATGATGACAAGAAGTAGC |
| | TM_FP3 | CGAAAAATGAAAAGGTCTTCTTCTGAAATAAAGACTGACTACC |
| | TM_RP3 | GGTGGCGACCGGTGGATCCACGATGGGTGGGGTGGAGTACAGGA |
| VEGFR1-JM$^{VEGFR2}$ | JM_FP1 | TACCGGACTCAGATCTCGAGATGGTCAGCTACTGGGACACCGG |
| | JM_RP1 | TTGGCCCGCTTAACGGTCCGGATAAAGAGGGTTAATAGGAGCC |
| | JM_FP2 | CGG ACC GTT AAG CGG GCC |
| | JM_RP2 | CCG GTC TCT GGG GAA TTC C |
| | JM_FP3 | GGGAATTCCCCAGAGACCGGCTTAAACTGGGCAAATCACTTG |
| | JM_RP3 | GGTGGCGACCGGTGGATCCACGATGGGTGGGGTGGAGTACAGGA |
| VEGFR1-TMJM$^{VEGFR2}$ | TMJM_FP1 | TACCGGACTCAGATCTCGAGATGGTCAGCTACTGGGACACCGG |
| | TMJM_RP1 | AGAATAATGATTTCCAAGTTAGACTTGTCCGAGGTTCCTTGAACAG |
| | TMJM_FP2 | AACTTGGAAATCATTATTCTAGTAGGCACGGC |
| | TMJM_RP2 | CCGGTCTCTGGGGAATTCCCATTTGC |
| | TMJM_FP3 | GGGAATTCCCCAGAGACCGGCTTAAACTGGGCAAATCACTTGGAAGAGGGG |
| | TMJM_RP3 | GGTGGCGACCGGTGGATCCACGATGGGTGGGGTGGAGTACAGGA |
| VEGFR1 ΔSSS | Fwd | CCCTCTTTATCCGAAAAATGAAAAGGGAAATAAAGACTGACTACCTATC |
| | Rev | GATAGGTAGTCAGTCTTTATTTCCCTTTTCATTTTTCGGATAAAGAGGG |
| VEGFR1-K1079Q | Fwd | GACAAAATCTACAGCACCCAGAGCGACGTGTGG |
| | Rev | CCACACGTCGCTCTGGGTGCTGTAGATTTTGTC |
| | Fwd | CTGGCACAGAGACCCATCAGAAAGGCCAAGATTTGC |

| | | |
|---|---|---|
| VEGFR1-R1142S | Rev | GCAAATCTTGGCCTTTCTGATGGGTCTCTGTGCCAG |
| VEGFR1-R1146T | Fwd | CCATCAGAAAGGCCAACATTTGCAGAACTTGTGG |
| | Rev | CCACAAGTTCTGCAAATGTTGGCCTTTCTGATGG |
| VEGFR1 -D1022N | Fwd | GTG CAT TCA TCG GAA CCT GGC AGC GAG |
| | Rev | CTCGCTGCCAGGTTCCGATGAATGCAC |
| HA-VEGFR1 | Fwd | CCTGAACTGAGTTTAAAAGGCACCC |
| | Rev | AGCGTAATCTGGAACATCGTATGGGTAATCTTTTAATTTTGAACCTGAACTAGATCC |
| HA-VEGFR1 ΔECD | Fwd | GTTCAAGGAACCTCGGACAAGTC |
| | Rev | AGCGTAATCTGGAACATCGTATGGGTAATCTTTTAATTTTGAACCTGAACTAGATCC |
| HA-VEGFR2 ΔECD | Fwd | GAAGGTGCCCAGGAAAAGACGAAC |
| | Rev | AGCCTTAATTGTAAGTATGTCTTTTTGTATGC |
| HA-VEGFR1 ΔECD -D1022N | Fwd | GTG CAT TCA TCG GAA CCT GGC AGC GAG |
| | Rev | CTCGCTGCCAGGTTCCGATGAATGCAC |
| HA-VEGFR1-TM$^{VEGFR2}$ ΔECD | Fwd | GTG CAT TCA TCG GAA CCT GGC AGC GAG |
| | Rev | CTCGCTGCCAGGTTCCGATGAATGCAC |
| HA-VEGFR1-TMJM$^{VEGFR2}$ ΔECD | Fwd | GTG CAT TCA TCG GAA CCT GGC AGC GAG |
| | Rev | CTCGCTGCCAGGTTCCGATGAATGCAC |
| HA-VEGFR1-JM$^{VEGFR2}$ ΔECD | Fwd | GTG CAT TCA TCG GAA CCT GGC AGC GAG |
| | Rev | CTCGCTGCCAGGTTCCGATGAATGCAC |
| VEGFR1-TM$^{gPA}$- G83I | Fwd | GTGATGGCTATTGTTATTGGAACG |
| | Rev | CCCAAAAATAATGAGCTCCAG |
| VEGFR1-TM$^{gPA}$ | gpA_FP1 | TACCGGACTCAGATCTCGAGATGGTCAGCTACTGGGACACCGG |
| | gpA_RP1 | CTCCAGATTAGACTTGTCCGAG |
| | gpA_FP2 | CGGACAAGTCTAATCTGGAGCTCATTATTTTTGGGGTGATGG |
| | gpA_RP2 | ACCGTAAGAAATTAAGAGGATCGTTCCAATAACACC |
| | gpA_FP3 | TCCTCTTAATTTCTTACGGTATC CGA AAA ATG AAA AGG TCT TC |
| | gpA_RP3 | GGTGGCGACCGGTGGATCCACGATGGGTGGGGTGGAGTACAGGA |
| | Fwd | CTG GAG CTG ATC ATT CTA ACA TGC ACC |

| | | |
|---|---|---|
| T667V | Rev | GGT GCA TGT TAG AAT GAT CAG CTC CAG |
| T669I | Fwd | GAGCTGATCACTCTAGTATGCACCTGTGTGGC |
| | Rev | GCCACACAGGTGCATACTAGAGTGATCAGCTCC |
| C670G | Fwd | CTGATCACTCTAACAGGCACCTGTGTGGCTG |
| | Rev | CGCAGCCACACAGGTGCCTGTTAGAGTGATCAGC |

**Table-S3**: Diffusion coefficient of different VEGFR1 and VEGFR2 constructs

| Constructs | $D_{confocal}$ ($\mu m^2.s^{-1}$) | |
|---|---|---|
| | -VEGF$_{165}$ | +VEGF$_{165}$ |
| 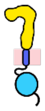 VEGFR1- wt | 0.0385 ± 0.0179 | 0.0177 ± 0.0071 |
| 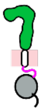 VEGFR2-wt | 0.0206 ± 0.0075 | 0.0107 ± 0.0040 |
| 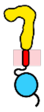 VEGFR1-TM$^{GPA}$ | 0.0211502 ± 0.0054 | - |
| 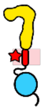 VEGFR1-TM$^{GPA(G83I)}$ | 0.0332 ± 0.0125 | - |
| 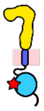 VEGFR1-D1022N | 0.0325 ± 0.01509 | 0.01565 ± 0.0056 |
| 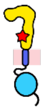 VEGFR1-C471R | 0.0348 ± 0.0142 | 0.0172 ± 0.0071 |
| 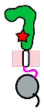 VEGFR2-C482R | 0.0176 ± 0.0060 | 0.0125 ± 0.0025 |
| 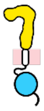 VEGFR1-TM$^{VEGFR2}$ | 0.022353 ± 0.0068 | 0.0129 ± 0.0032 |
| 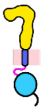 VEGFR1-JM$^{VEGFR2}$ | 0.0205 ± 0.0059 | 0.0127 ± 0.0047 |
| 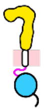 VEGFR1-TMJM$^{VEGFR2}$ | 0.0202 ± 0.0070 | 0.0131 ± 0.0050 |

**Table-S4**: Tyrosine Phosphorylation rate and phosphorylation half-life of VEGFR constructs after treatment with $VEGF_{165}$ or Sodium orthovanadate mediated

| | Construct | Phosphorylation rate (A.U. $min^{-1}$) | | Phosphorylation half-life (min) | |
|---|---|---|---|---|---|
| | | $+VEGF_{165}$ | $+Na_3VO_4$ | $VEGF_{165}$ mediated | $Na_3VO_4$ mediated |
| 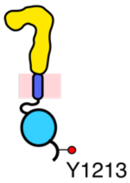  | VEGFR1-wt | 0.07 ± 0.01 | 0.025 ± 0.003 | 14.3 ± 4.4 | 12.0 ± 1.6 |
| 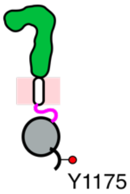  | VEGFR2-wt | 0.17 ± 0.02 | 0.051 ± 0.005 | > 60 | 41.4 ± 4.2 |
| 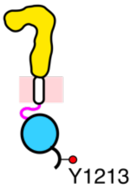  | VEGFR1-TM-$JM^{VEGFR2}$ | 0.16 ± 0.03 | 0.048 ± 0.008 | 48.6 ± 13.8 | 30.3 ± 7.3 |

**Table S5**: RMSDs of VEGFR1 model structures with respect to VEGFR1 crystal structure (PDB ID: 3HNG).

| Top 5 Models of VEGFR1 | Backbone RMSD wrt. VEGFR1 (PDB ID: 3HNG) (in Å) |
|---|---|
| Model 1 | 0.589 |
| Model 2 | 0.617 |
| Model 3 | 0.569 |
| Model 4 | 0.599 |
| Model 5 | 0.822 |

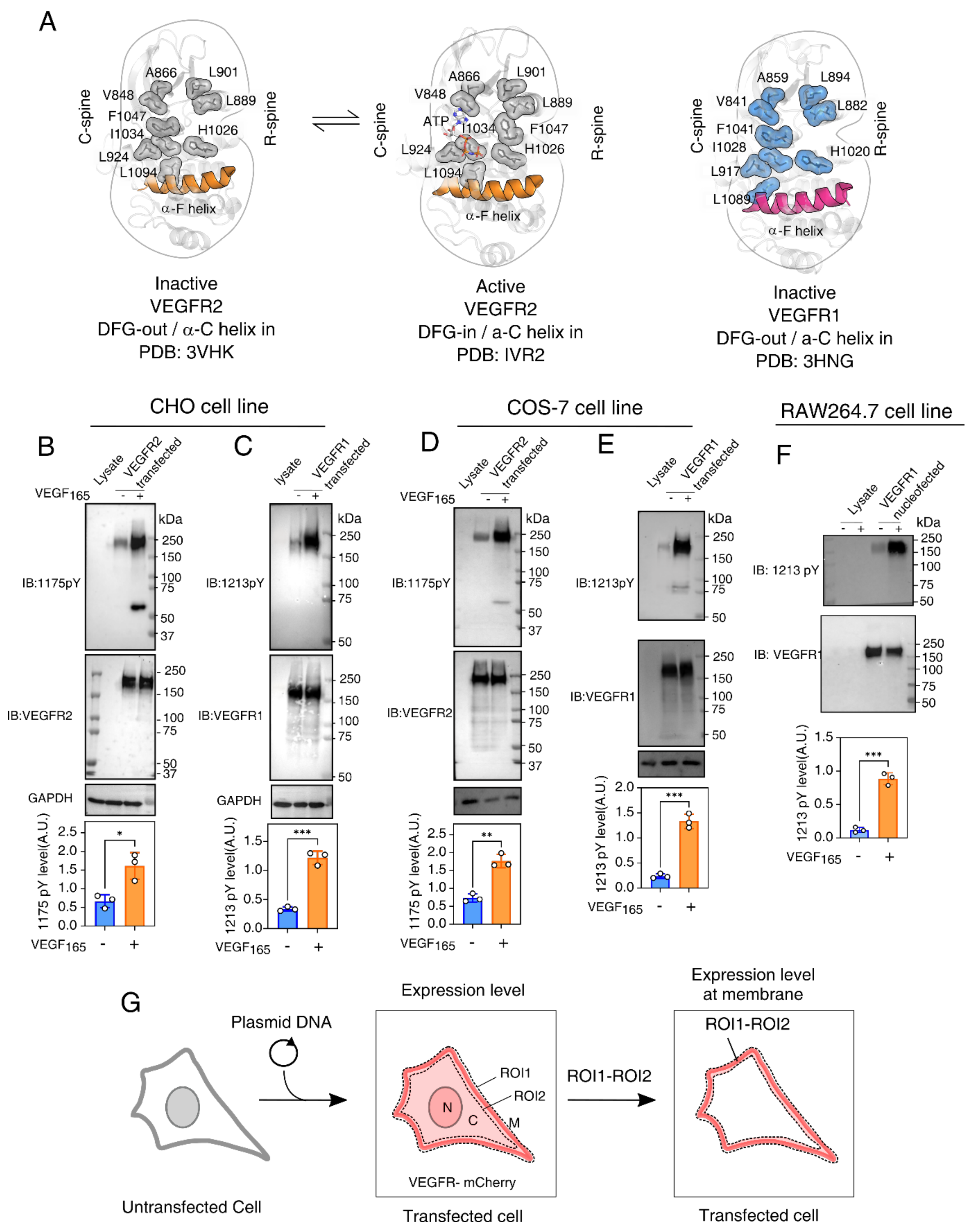


**Figure S1. Activation of VEGFR1 in comparison to VEGFR2 in COS7 and RAW264.7 cell line.**

**(A)** Structural analysis of conserved signature motifs (Regulatory and Catalytic spine) in the indicated crystaVEGFR2 and VEGFR1 kinase domain crystal structures.

**(B-F)** Representative immunoblot determining the tyrosine phosphorylation level of indicated C-terminal tyrosine residue of VEGFR2 (panels B and D) and VEGFR1 (panels C, E, and F), respectively. The VEGFR constructs were transiently expressed in the indicated cell type and activated by treating with $VEGF_{165}$. Bar plots in the lower panel are the densitometric quantitation of the Y1175 (panels B and D) or Y1213 (panels C, E, and F) phosphotyrosine levels from the respective immunoblots.

**(G)** Schematic representation of the single-cell assay of VEGFR activation. The area covered by ROI1 and ROI2 is shown by dotted lines. The membrane fraction is obtained by subtracting ROI2 from ROI1.

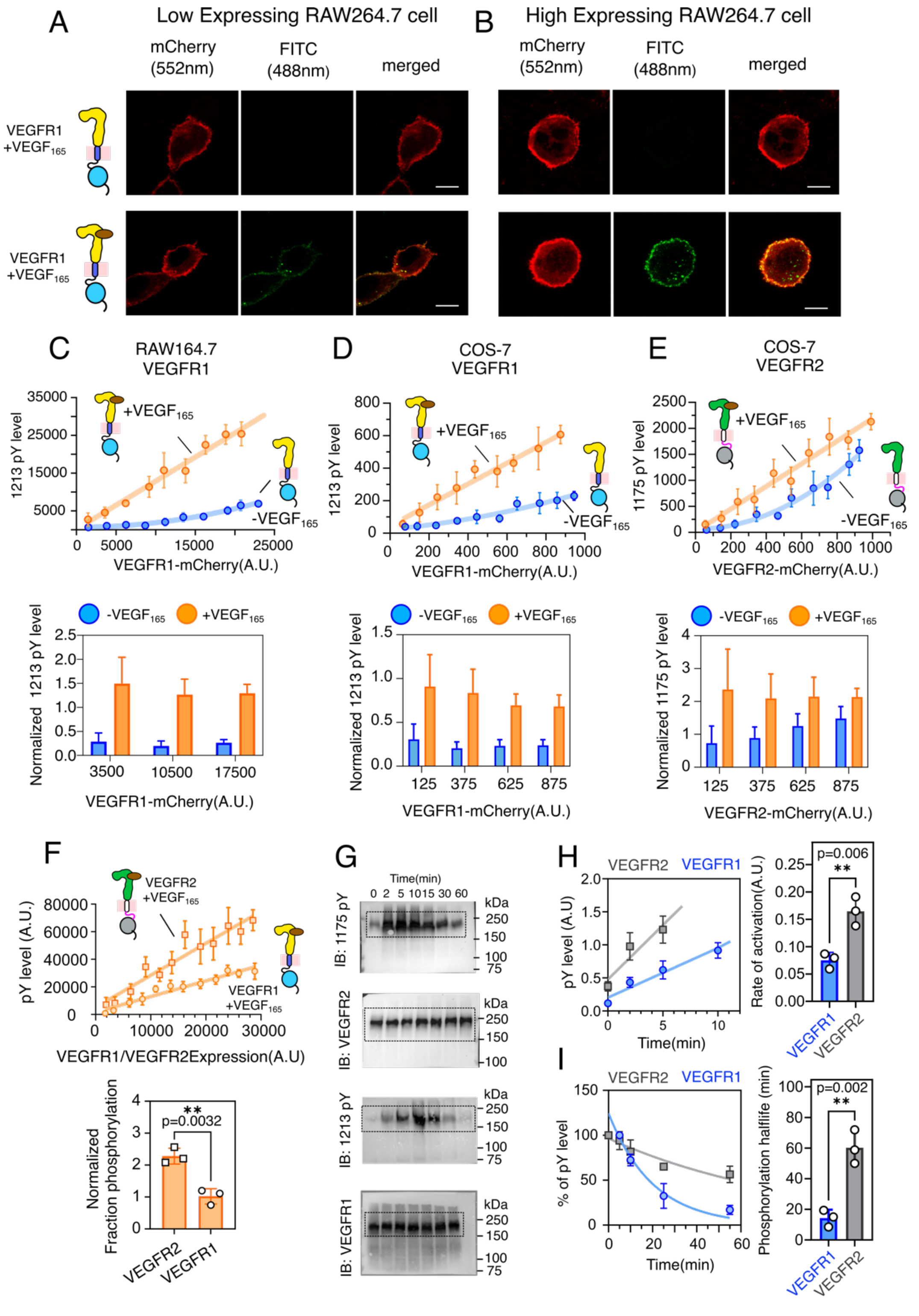


**Figure S2: Single-cell analysis of VEGFR activation and tyrosine phosphorylation kinetics**

**(A-B)** Confocal images of VEGFR1 fused to mCherry and Y1213 phosphorylation level (in green) were measured in a transiently RAW264.7 cell line. Panels A and B show the phosphorylation level of Y1213 in low or high expressing VEGFR1, respectively, with and without ligand treatment. Scale bar = 5 µm

**(C-D)** The Y1213 phosphorylation and VEGFR1 expression level in RAW264.7 and COS-7 cell lines, respectively, are plotted. Individual data points in the upper panel represent the mean expression and phosphorylation level for the selected cell with comparable expression levels (2500 A.U.). Individual data points for activation in the presence of $VEGF_{165}$ were fitted to a linear equation. In ligand-independent conditions, the blue lines show the second-order polynomial fit. In the lower panel, the bar graphs depict the mean phosphorylation ratio to its expression level within an expression range of comparable magnitude. The error bar shows the standard deviation of data points.

**(E)** The Y1175 phosphorylation is plotted against the expression level of VEGFR2 transiently expressed in COS-7 cell line. The bar graphs, in the lower panel, reflect the mean phosphorylation ratio to its expression level within an expression range of similar magnitude. The error bar shows the standard deviation of data points.

**(F)** The upper panel compares the ligand-dependent phosphorylation levels of VEGFR1 and VEGFR2 as a function of their expression levels. The bar plot at the bottom depicts the fraction of VEGFR1 or VEGFR2 phosphorylation, as determined from the slope of the curve fitted to a linear equation.

**(G)** Representative uncropped blots of the main Figure 2(G)

**(H)** The left panel shows the fitting of the densitometric analysis of Y1175 (grey) and Y1213 (blue) phosphorylation as a function of time in Figure 2G to a linear equation. The bar plot at right shows the rate of phosphorylation as determined by the slope.

**(I)** The left panel plots the densitometric analysis of Y1175 (gray) and Y1213 (blue) dephosphorylation as a function of time in Figure 2G to a linear equation. The bar plot shows the dephosphorylation rate as determined from fitting the data points to the exponential decay curve (left panel).

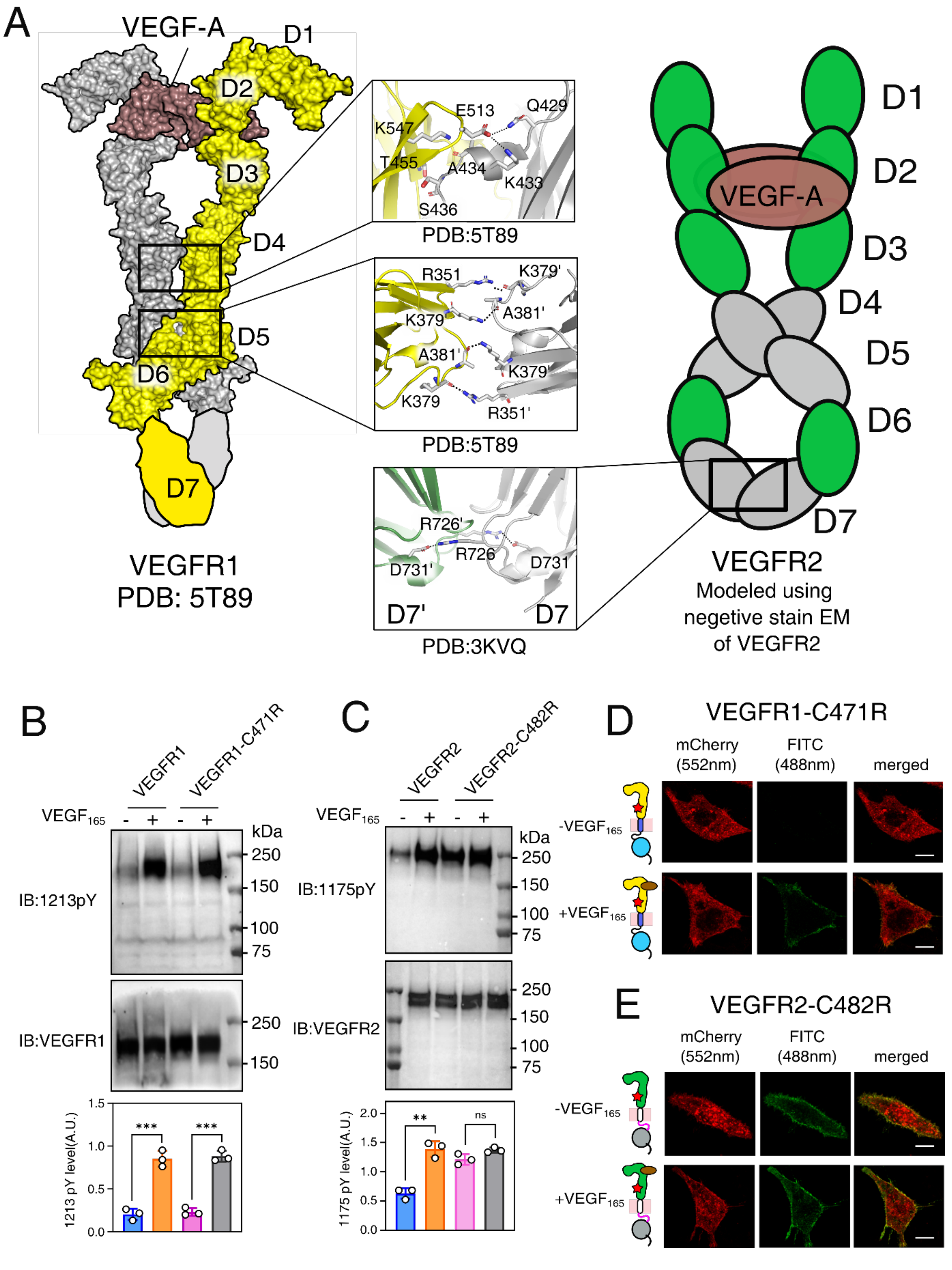


**Figure S3: Function of ECD and constitutively active mutant in ligand-dependent VEGFR activation**

**(A)** Left panel shows the VEGFR1(D1-D6) cryo-EM structure. The D7 is modeled using VEGFR2-D7 as a template[19]. The right panel shows a schematic model of VEGFR2 (D1-D7) from a negative stain electron micrograph [20]. The middle panel shows the cross-section of the dimerization arms (D4, D5, and D7). The key residues at the dimer interface are labelled.

**(B)** In the upper panel, Immunoblot shows the Y1213 phosphorylation of VEGFR1-C471R mutant after with and without ligand treatment. Bar plots in the lower panel represent the densitometric quantitation of the Y1213 phosphorylation level.

**(C)** In the upper panel, immunoblot shows the Y1175 phosphorylation of constitutively activating VEGFR2-C482R mutant in the presence and absence of ligand. Bar plots in the lower panel represent the densitometric quantitation of the Y1175 phosphorylation level.

**(D-E)** Confocal images showing the phosphorylation level of mCherry fused VEGFR1-C471R (D) or VEGFR2-C482R (E) in transiently transfected CHO cell lines.

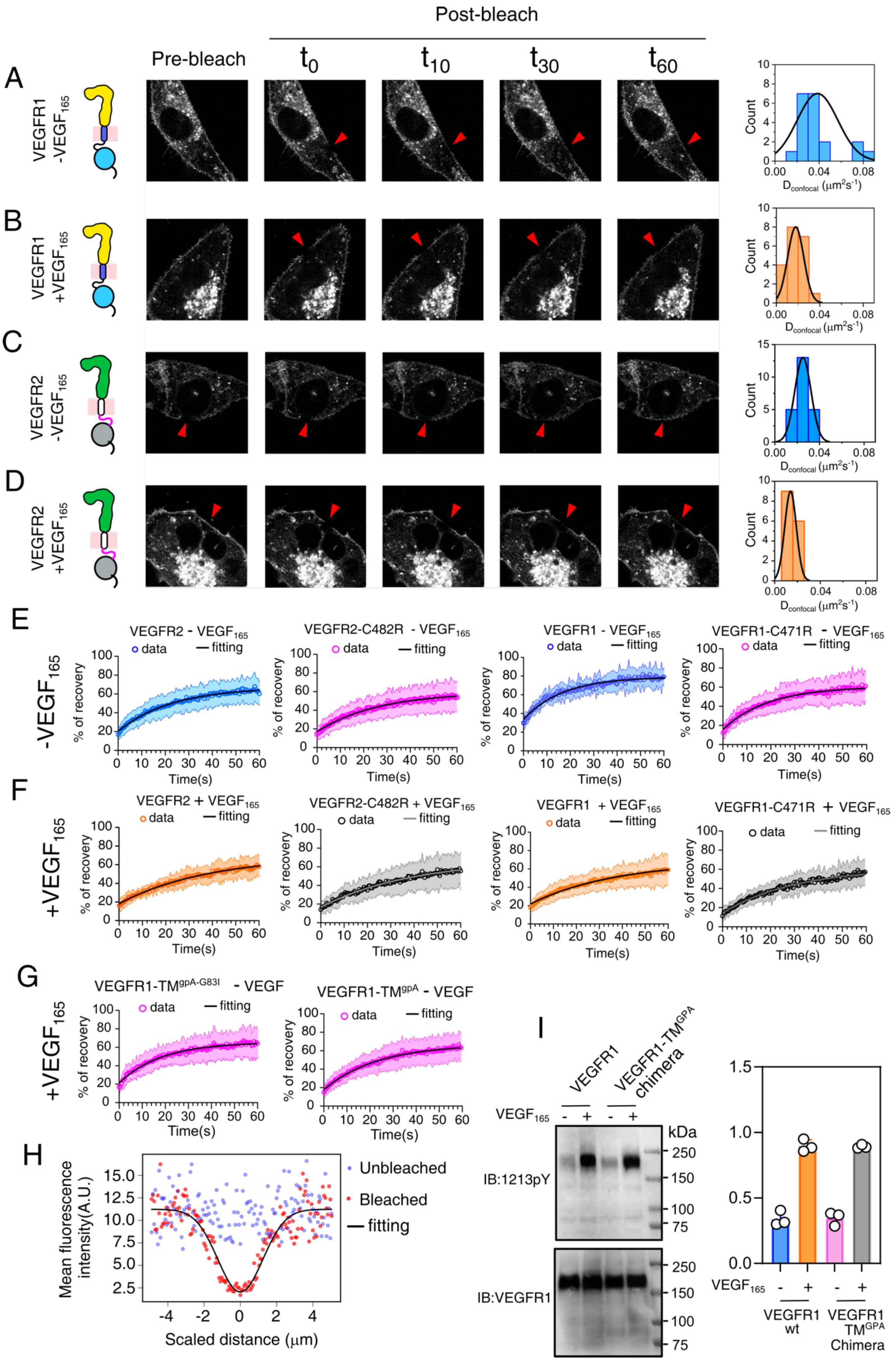


**Figure S4: Analysis of VEGFR dimer by FRAP**

**(A-D)** Time-lapse image of CHO cell line transiently expressing indicated constructs of VEGFR. FRAP was performed on the plasma membrane of VEGFR1 (panels A, and B) or VEGFR2 (panels C, and D) transfected CHO cell line in the presence(panels B, and D) or absence (panels A, and C) of $VEGF_{165}$. A red arrow indicates the bleach spot. Recovery of fluorescence is shown at indicated time points. The right panel shows the normal distribution of diffusion coefficients. Number of cells 25-30

**(E-F)** FRAP profile of indicated constructs in the absence (E) or presence (F) $VEGF_{165}$ , each profile is fitted to the first order exponential equation.

**(G)** FRAP profile of monomer (left panel) and dimer(right panel) control, as explained in Figure 3, in the absence of $VEGF_{165}$.

**(H)** Determination of effective bleach spot radius($r_e$) from post-bleach intensity profile.

**(I)** Immunoblot (left panel) showing the Y1213 phosphorylation level of VEGFR1-$TM^{GPA}$ (TM dimer control) as compared to its wild-type control. Bar plots in the right panel represent the quantitation of the Y1213 phosphorylation level.

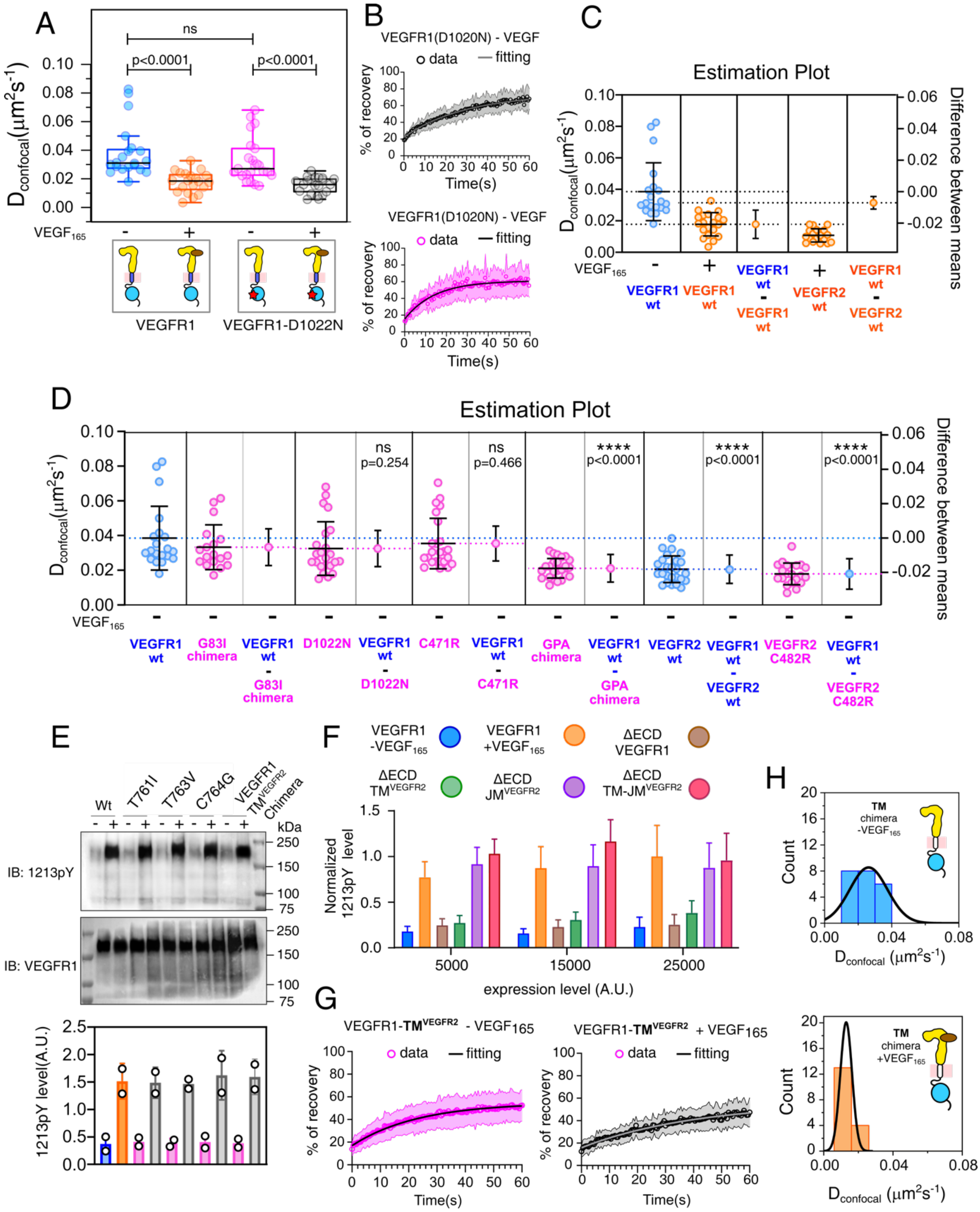

**Figure S5: Study of TM segment in activating VEGFR1**

**(A)** Box plot of the Diffusion coefficient of VEGFR1 and kinase-dead mutant (D1022N) determined from FRAP experiment. Each data point in the box plot reflects the diffusion coefficient of the selected cell, while the black line indicates the mean value. Number of cells 20-30

**(B)** FRAP profile of VEGFR1 kinase-dead mutant (D1022N) in the presence (upper panel) and absence (bottom panel) of $VEGF_{165}$.

**(C-D)** The estimation plot shows the mean difference between the two indicated data sets. The left axis indicates the values of the diffusion coefficient, and the right axis is the difference between the mean at the 95% confidence range.

**(E)** Top panel is the Immunoblot of Y1213 phosphorylation in transmembrane domain mutants and chimeric constructs of VEGFR1. The densitometric quantification of Y1213Y phosphorylation is depicted by bar graphs (bottom panel).

**(F)** The bar plots representing the normalized Y1213 phosphorylation level of VEGFR1-ECD deleted constructs, shown in Figure 4C.

**(G)** FRAP profile of VEGFR1 -$TM^{VEGFR2}$ chimera in the presence and absence of $VEGF_{165}$.

**(H)** Normal distribution of diffusion coefficient measured for VEGFR1 -$TM^{VEGFR2}$ chimera in the presence and absence of ligand.

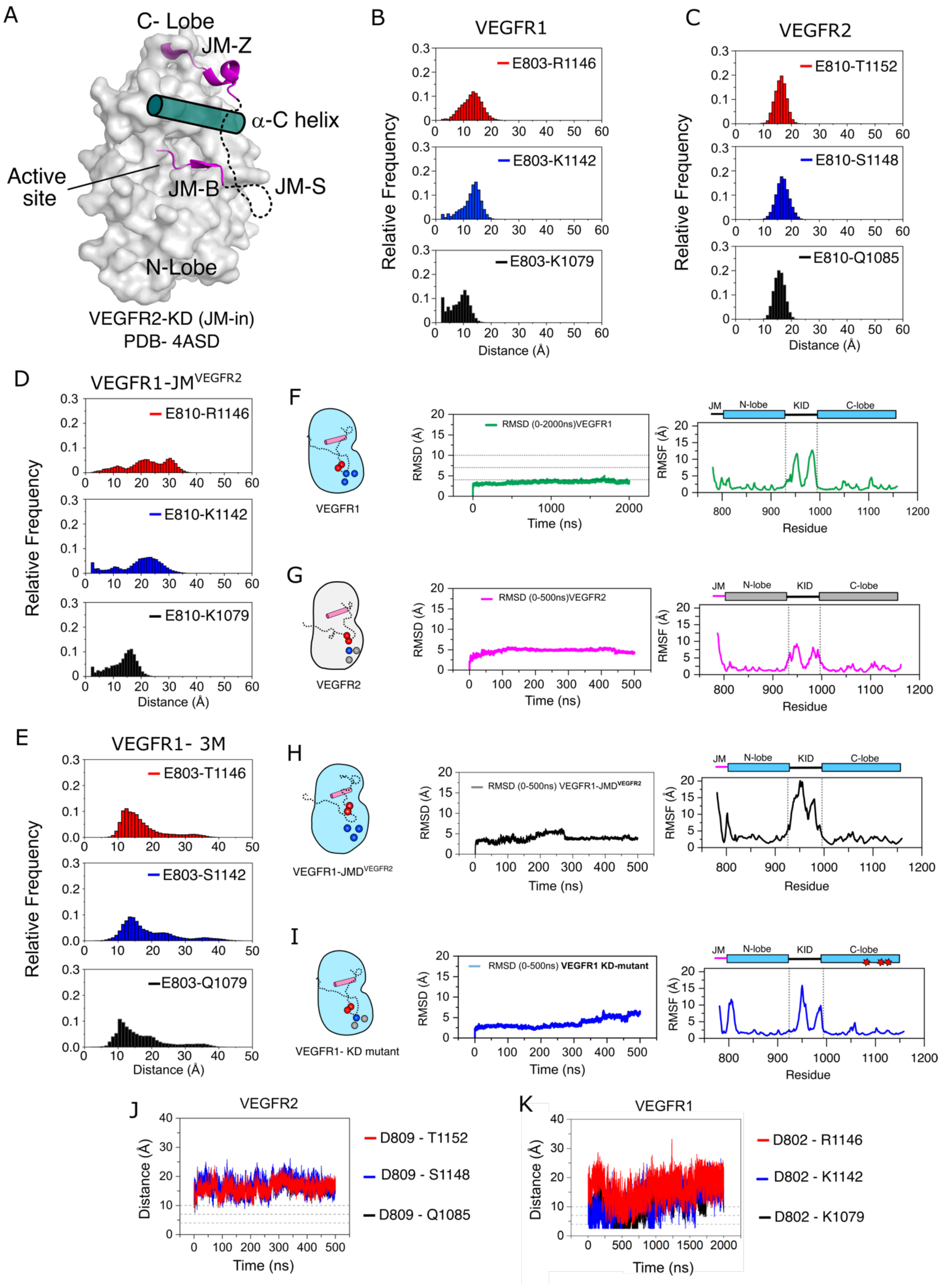


**Figure S6: Molecular dynamics simulation of JM-KD of VEGFR**

**(A)** Inactive structure of VEGFR2 in JM-in conformation is shown as a space-filled model. The JM-S segment (in dotted lines) is modeled based on the PDGFR crystal structure (PDB ID: 5K5X).

**(B-E)** Population distribution of indicated ion pairs in the electrostatic latch for the construct described above (panel B-E).

**(F-I)** The root mean square deviation (RMSD) and root mean square fluctuation (RMSF) for the indicated JM-KD constructs of VEGFR.

**(J-K)** The pairwise interatomic distances determined for the residues in the electrostatic latch are plotted against time for the constructs described in Figures 5 B and C, respectively. The horizontal lines represent the distance cut-off for the weak, medium, and strong electrostatic interactions [21, 22].

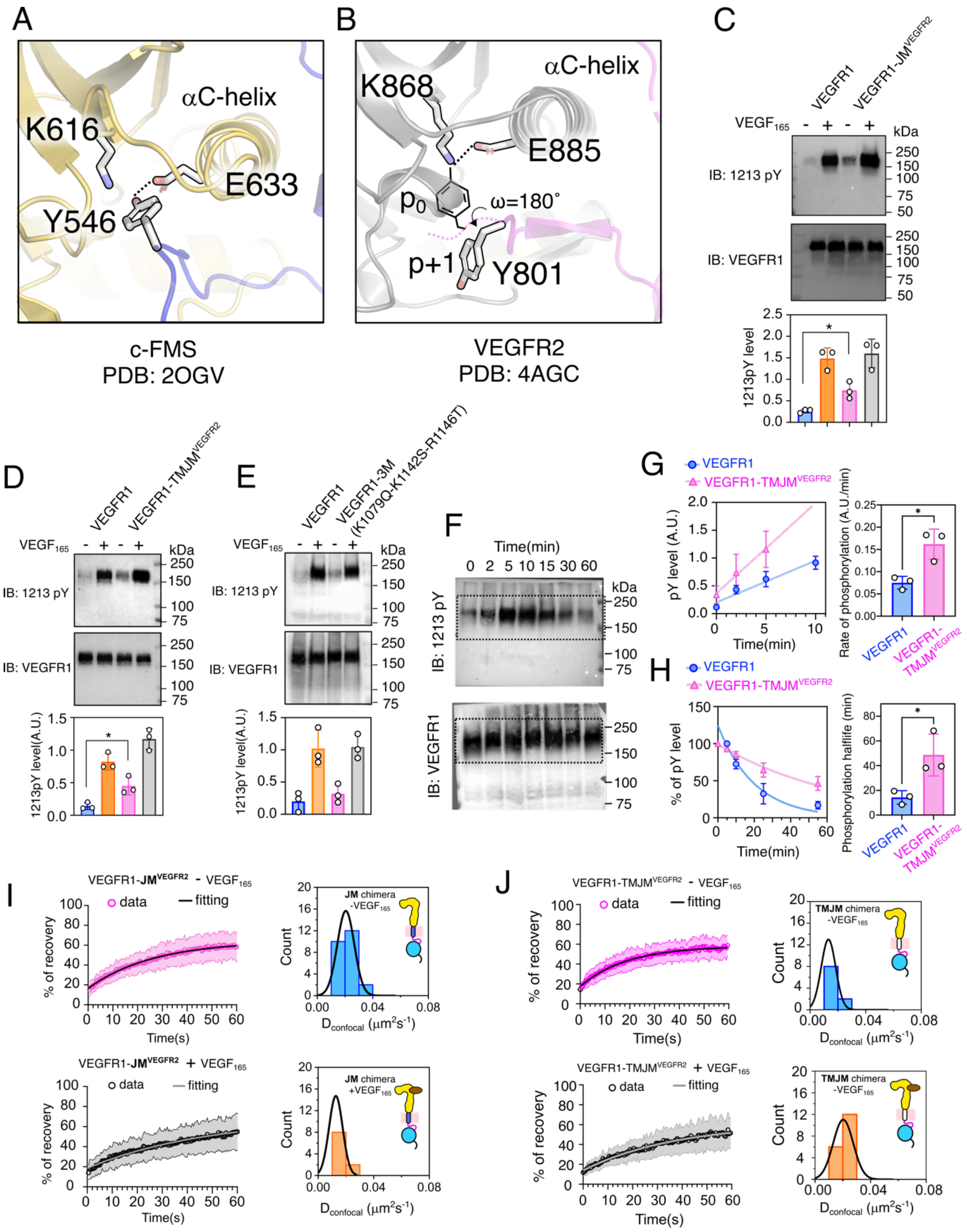


**Figure S7: Study of JM inhibition**

**(A)** A close-up view of the inhibitory interaction between the Tyr546 and Glu633 in the JM-B and C-helix of the cFMS KD crystal structure (PDB ID: 2OGV).

**(B)** Close-up view of the autoinhibitory conformation of the JM-B segment in the VEGFR2 crystal structure. The orientation of the conserved Y801 residue in the JM-B is shown. The expected orientation of the Y801 residue is drawn considering the Y801 residue is moved to the -1 position, and the ω angle is rotated by 180°.

**(C-E)** The upper panel shows the immunoblot of Y1213 phosphorylation in the indicated VEGFR1 constructs. Bar plots in the bottom panel represent the densitometric quantification of the Y1213 phosphorylation level from three independent experiments.

**(F)** Representative uncropped blots shown in main Figure 6 D

**(G)** The rate of Y1213 phosphorylation is determined from the slope of the phosphorylation level vs time plot, measured from the densitometric analysis of Figure 6D. The phosphorylation rate is determined from the linear fit of phosphorylation at time $t_0$ to the highest phosphorylation level measured. The right panel shows the average phosphorylation rates measured from three independent experiments.

**(H)** The half-life of the phosphotyrosine residue Y1213 of VEGFR1 is determined from the exponential fitting of the phosphorylation decay from the highest phosphorylation observed in Figure 6D. The right panel shows the average phosphorylation half-life measured from three independent experiments.

**(I-J)** The left panel shows the FRAP profile of VEGFR1 -JM$^{VEGFR2}$ and VEGFR1 -TM-JM$^{VEGFR2}$ chimeras, respectively, in the presence and absence of ligand. The solid-line represents the fitting to a first-order exponential equation.The right panel represents the normal distribution of the diffusion coefficient of indicated constructs.

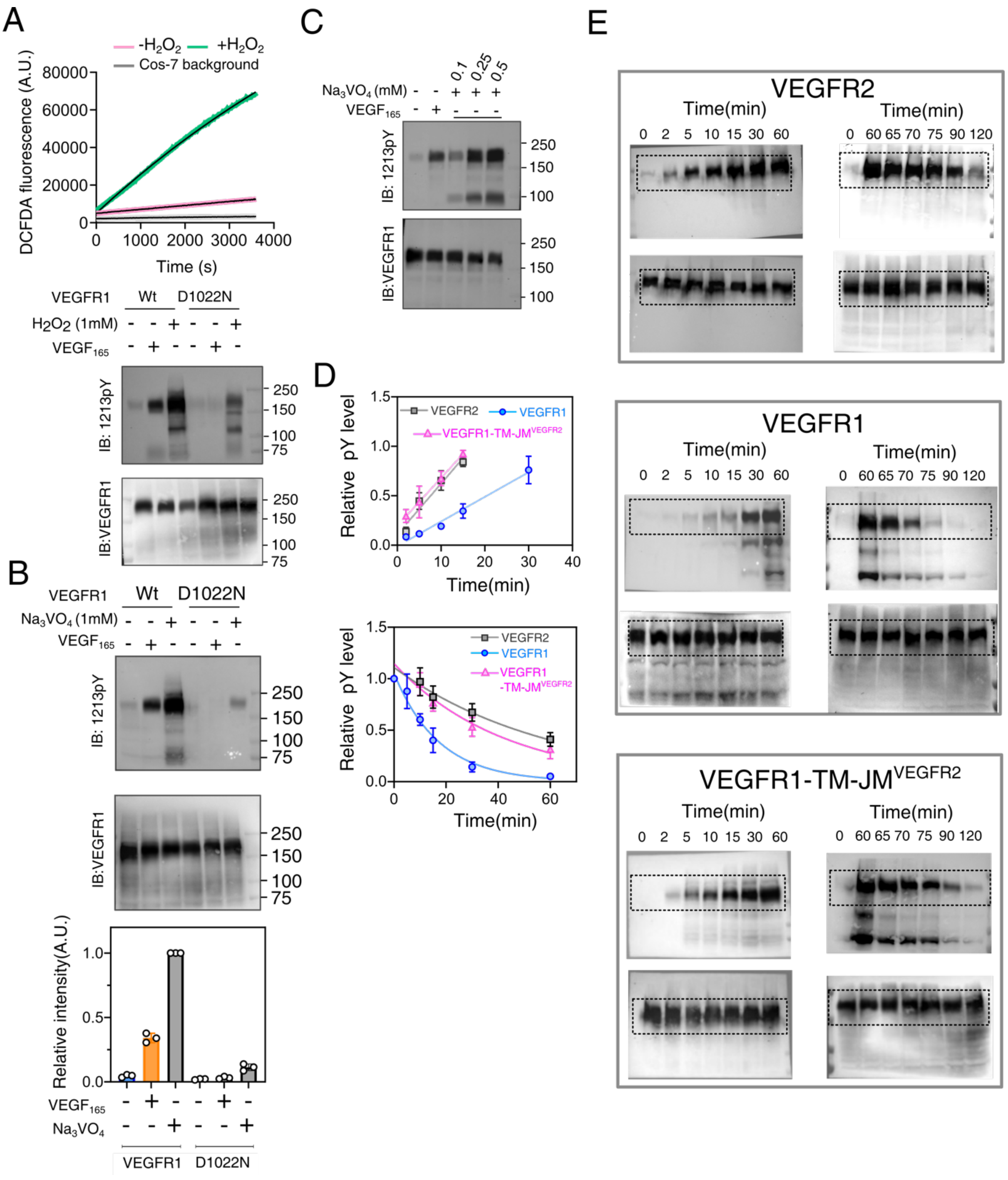


**Figure S8: Inhibition of PTP results in increased ligand-independent autophosphorylation of VEGFR1**

**(A)** ROS generation in the Cos-7 cell line is detected by measuring the change in DCFDA fluorescence over one hour after treating cells with $H_2O_2$. The bottom panel represents the immunoblot of VEGFR1 Y1213 phosphorylation after treatment with $H_2O_2$.

**(B)** Representative immunoblot of VEGFR1 Y1213 phosphorylation level compared to a kinase-dead mutant (D1022N) at the indicated experimental condition. Below, is the densitometric analysis of Y1213 phosphorylation level for the indicated VEGFR1 constructs.

**(C)** Representative immunoblot determining the effect of increasing $Na_3VO_4$ concentration on ligand independent-phosphorylation of Y1213.

**(D)** Individual data represents the tyrosine phosphorylation level at the indicated time point is determined from the densitometric analysis of immunoblots shown in Figures 7C, and D. Data sets were fitted to a linear (Figure

7E) and exponential decay (Figure 7E) function for measuring the phosphorylation rate and half-life, respectively.

**(E)** Representative uncropped blots shown in main figure 7(C-D)
